# The Divergent 3D Genome Landscapes of Aging and Neurodegenerative mouse models

**DOI:** 10.1101/2025.11.10.687599

**Authors:** Ekaterina Kashuk, Dmitrii Smirnov, Ekaterina Eremenko, Alexandra Tsitrina, Dmitrii Kriukov, Anastasia Golova, Shai Kaluski-Kopatch, Ekaterina Khrameeva, Debra Toiber

**Affiliations:** Center for Molecular and Cellular Biology, Skolkovo Institute of Science and Technology, Moscow 121205, Russia; Department of Life Sciences, Ben-Gurion University of the Negev, Beer Sheva 8410501, Israel; The Zlotowski Center for Neuroscience, Ben-Gurion University of the Negev, Beer Sheva 8410501, Israel; Ilse Katz Institute for Nanoscale Science and Technology, Ben-Gurion University of the Negev, Beer Sheva 8410501, Israel

## Abstract

Chromatin structure is essential for gene regulation and genome stability, and neurons must preserve their 3D genome organization throughout life. This structure gradually deteriorates with aging and is further disrupted in neurodegenerative diseases. To investigate whether aging and neurodegeneration share early chromatin changes or diverge, we performed Hi-C on cortical neurons from adult, aged, and brain-specific SIRT6-knockout (S6-KO) mice, and compared them to the CK-p25 Alzheimer’s disease model results stratified by γH2AX levels. All models showed early features such as weakened interactions between chromosomes in expanded nuclei, A-to-B compartment shifts, and loss of architectural loops. However, aged neurons retained TAD prominence, short-range interactions, and enhancer-promoter loops, while pathological models showed reduced TAD prominence, shorter loops, and disorganized A–B mixing. These changes correlate with increased DNA damage. Our findings suggest that aging represents a “primed” state, where chromatin regulation begins to erode, but further stressors like DNA damage are associated with progression toward neurodegenerative breakdown.

## Introduction

Aging is the main cause of neurodegeneration, and DNA damage accumulation has been considered the main driver of aging and disease. However, the question of whether natural aging is an earlier stage before dementia or two separate pathways could be relevant for decisions on how to prevent and treat neurodegenerative diseases.

Sporadic cases of AD incidence may arise from very different causes, such as metabolic syndromes, head injury, depression and more. And although these are different factors, however, at some points neuronal stress responses, such as overexpression of Tau, loss of proteostasis, DNA damage response and cell death, are all activated, producing similar symptoms, making all fall under the umbrella of Alzheimer’s disease.

Various animal models of neurodegeneration have been developed, among them SIRT6 deficiency is an important model^1,2,3,4^. SIRT6 is a chromatin bound protein. One of its most important activities is DNA damage repair^5,6^, where it acts as a DNA damage sensor, allowing chromatin accessibility and recruiting repair and restoration proteins to the sites of damage. Moreover, through its catalytic activities, SIRT6 regulates gene expression, metabolic homeostasis, inflammation, genome maintenance, and lifespan^7,8,9^.

Another important model is the CK-p25 model, which is based on the overexpression of the p25 protein, a cleavage product of p35. The accumulation of p25 aberrantly activates Cyclin-Dependent Kinase 5 (Cdk5) and Glycogen Synthase Kinase-3b (GSK3b). This dysregulated kinase activity leads to the hyperphosphorylation of tau, synaptic dysfunction, and neuronal death. The model can also show elevated levels of amyloid-β and synaptic deficits, further linking it to AD-like pathology.

Although these models initiate different pathways, both present DNA damage, cognitive impairment and learning disabilities, increased apoptosis in the brain, accumulation of hyperphosphorylated Tau, and reduced genomic stability, together resulting in a neurodegeneration-like phenotype, similar to the different “causes” in AD having a similar outcome in the brain.

During aging cortical neurons present a non-senescent nuclei expand, leading to increased distances between chromosomes, affecting the topology of compartments, topologically associating domains (TADs) and chromatin loops. These topological changes impact the borders of TADs, resulting in border weakening. Moreover, this expansion leads to nuclear envelope weakening and deformations^10^. Chromatin landscape is very important to assure cellular differentiation and maintenance, we and others have observed critical changes in chromatin organization during senescence^11,12,13^, aging^10,14,15,16,17,18^ and in diseases^19,20,21,22^.

However, these mice experience a healthy aging, and whether this aged-state is similar or a precursor to neurodegeneration was not clear. In order to better understand neurodegeneration we decided to focus on SIRT6 as an pathological aging accelerated model. Notably, SIRT6 levels were shown to decrease with age, which multiple research groups have speculated to promote the aging process^23,24,1,25^. Furthermore, when examining SIRT6 levels in the brains of Alzheimer’s disease patients, they exhibit much lower SIRT6 expression compared to healthy aged individuals^1^. In addition to the CK-p25 model, allowing us to compare two different initiators of disease that culminate in very similar phenotypes.

In this work we show that chromatin organization in neurons is dynamically changing during aging and SIRT6 deficiency, however, during pathological changes, significant divergence occur, particularly TAD prominences loss is only observed in late stages, while loss of inter-chromosomal interactions and weaker boundaries are common also to normal aging. Importantly, these changes are correlated with the increase in DNA damage, occurring by the lack of SIRT6 and increase in H2AX in the CK-p25 AD mouse model. Moreover, different transcription factors are associated with regular aging vs pathological aging, suggesting these biomarkers could be misregulated in pathological aging brains.

## Methods

### Primary Cortical Neuron Culture

Primary cortical cultures were established through the isolation of cortical neurons derived from pregnant C57BL/6 mice. To summarize, cortical tissues were harvested and underwent dissociation facilitated by papain (Sigma-Aldrich, P3125). Subsequently, the dissociated neurons were seeded onto poly-L-lysine-coated chambers (μ-slide 4-well glass bottom, IBIDI GmbH Martinsried, Germany, 80427) utilizing Neurobasal medium (GIBCO, 21103–049), supplemented with B-27 (GIBCO, 17504044), GlutaMAX (GIBCO, 35050–061), and 2% FBS (HyClone). Following a 24-hour incubation period, the medium was replaced with serum-free Neurobasal medium. Lentiviral infection was introduced to the neurons after 5 days. Cells were infected with lentivirus pLKO backbone system (Sigma-Aldrich). SIRT6 was targeted using the shRNA sequence CCGGGCATGTTTCGTATAAGTCCAACTCGAGTTGGACTTATACGAAACATGCTTTTTG, with scrambled shRNA used as a control. Five days after infection, cells were fixated with 4% Paraformaldehyde for 10 minutes.

### Generation of S6-KO mouse models

S6-KO mice were generated through Nestin-Cre-mediated conditional knockout as described in Etchegaray et al.^26^ Briefly, mice with loxP-flanked exon 2 of the SIRT6 gene were backcrossed for three generations and subsequently crossed with C57BL/Nestin-Cre/J transgenic mice to achieve Cre/LoxP-mediated excision of exon 2. Middle-aged (10-11 months old) S6-KO mice were selected for the subsequent analyses. All procedures involving animals were approved by the Institutional Animal Care and Use Committee (IACUC) at Massachusetts General Hospital and conducted in accordance with NIH guidelines.

### Tissue Collection and Nuclei Isolation

Nuclei isolation from the brain tissue and further NeuN antibody immunostaining were performed according to the previously published protocol^27^. Briefly, after isoflurane anesthesia and phosphate-buffered saline (PBS) perfusion, mouse cortical tissues were rapidly dissected on ice. The cortices were then homogenized in a 500 μl ice-cold low sucrose buffer and crosslinked with formaldehyde, followed by quenching with glycine. Following centrifugation, pellets were resuspended in 3 ml of low sucrose buffer supplemented with protease inhibitor cocktail (PIC) and homogenized. Nuclei were subsequently isolated via iodixanol gradient centrifugation, stained with NeuN-PE antibody, and sorted by fluorescence-activated cell sorting (FACS) after filtration.

### Generation of Hi-C libraries

Hi-C maps were generated for neuronal nuclei isolated using FANS from the cortex tissue of brain-specific SIRT6-knockout (S6-KO) adult (10-11 months) mice and control adult (10 months) and old (20 months) mice (3 mice per each group). Hi-C was performed using the DpnII restriction enzyme, according to a previously published protocol^28^. Briefly, approximately 1.2–1.4 million fixed neuron nuclei were resuspended in 100 μL total volume containing 1.1× DpnII buffer and 0.3% SDS, incubated for 1 h at 37°C with 1400 rpm shaking, followed by Triton X-100 quenching (1.8% final) for 1 h under identical conditions. DpnII restriction (200 U) was performed for 3–4 h (or 16–18 h) at 37°C with 1400 rpm shaking, then inactivated at 65°C for 20 min. After centrifugation (3200 g, 10 min, 20°C), nuclei were resuspended in 100 μL 1× NEB2 buffer. Biotinylation of DNA ends was conducted in 120 μL total volume for 1.5 h at 37°C with 1000 rpm shaking. Subsequent blunt-end ligation used 1000 U T4 ligase in 300 μL total volume for 16–18 h (or 5–6 h) at 20°C with 1400 rpm shaking. Reverse-crosslinking employed Proteinase K and SDS for 4–5 h (or 16–18 h) at 65°C. DNA was purified by phenol:chloroform and chloroform extractions, then ethanol-precipitated with sodium acetate and glycogen. After dissolution in Tris-HCl, DNA was sheared to <1000 bp by sonication (20 cycles, 30s ON/30s OFF, high power, 4°C). Size selection used Amicon columns (30-kDa) with Tris-HCl washes, followed by AmpureXP bead purification. Biotin pull-down employed MyOne Streptavidin C1 beads in low-binding tubes with Tween washing buffer (TWB) and binding buffer (BB) washes. Bead-bound DNA underwent end repair (30 min at 20°C, then 20 min at 75°C) and A-tailing (30 min at 20°C, then 20 min at 65°C). Adapter ligation used Illumina TruSeq adapters and Quick T4 ligase for 2 h at 20°C. After TWB and NEBuffer 2.1 washes, DNA was eluted from beads by heating (98°C, 10 min). Libraries were amplified by Real-Time PCR to determine optimal cycles, pooled, and purified with 1.8× AmpureXP beads (75% ethanol washes), then quantified fluorometrically and checked by agarose gel electrophoresis. Each Hi-C library was sequenced to 150-350 million paired-end reads.

### Preprocessing of Hi-C maps

The Hi-C sequencing reads were analyzed using the distiller software^29^ (https://github.com/open2c/distiller-nf, v.0.3.3). Reads were aligned to mm10 reference genome using *bwa* mapper^30^ and only those with a MAPQ greater than 30 were retained. Invalid read pairs were filtered out, and the remaining valid pairs were assigned to specific restriction fragments to construct a raw contact matrix. Raw contact Hi-C maps were downsampled to the same total number of contacts using the cooltools.sample function from the cooltools library v.0.5.1^31^. To minimize bias from Hi-C ligation artifacts, values along the main diagonal were excluded before sampling, which was performed using cooler and custom Python scripts. The resulting .cool files containing Hi-C contact matrices were then ICE normalized^32^ via the ‘cooler balance’ command with default settings. The obtained Hi-C maps binned from 1-kb to 1-Mb resolutions were further used as input for downstream analyses.

### Chromosomal scale analysis

The principal component analysis (PCA) was conducted based on the variation in the PC1 values obtained from all the generated Hi-C maps. Scaling curves were computed at 250-kb resolution by averaging the density of contacts relative to linear genomic distance. The analysis of interactions within all chromosomes (cis-contacts) and between all pairs of chromosomes (trans-contacts) was performed using in-house Python scripts deposited on GitHub:https://github.com/SIRT6/Sirt6_hic_paper. The compaction status of each 1-kb genomic region was assessed by determining its distal-to-local log_10_ ratio (DLR) and interchromosomal fraction (ICF) of Hi-C interactions at that specific locus. The DLR was calculated to determine the total number of interactions involving the 1 kb region of interest with regions located over 10.5 Mb upstream (defined as distal) compared to those located over 10.5 Mb downstream (defined as local). This threshold was established based on the intersection point of the scaling curves.

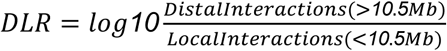

ICF was calculated by comparing the total number of Hi-C interactions that occurred between the 50-kb region of interest and other chromosomes to the total number of interactions at that specific locus.

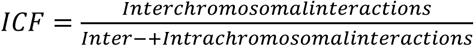

### Chromatin compartment analysis

Hi-C compartments were identified at a 50-kb resolution using *cooltools* software (v0.5.1)^31^ (https://zenodo.org/record/5214125) and the eigs_cis() function with a parameter setting of n_eigs = 8. The principal component (PC) showing the strongest correlation with the GC-content vector was used for further compartment analysis. Saddle plots were generated using the cooltools.saddle() function with the following parameters: contact_type=’cis’, n_bins=38, and qrange=(0.025, 0.975). The code for saddle plots was adapted from the cooltools tutorial: https://cooltools.readthedocs.io/en/latest/notebooks/compartments_and_saddles.html. For computing the strength of compartment interactions, the intensity of the top 10% of corresponding interactions derived from the saddle matrix was used. Compartment strength was calculated as follows:

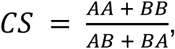

where AA, BB, AB and BA denote interaction intensities of the corresponding compartments extracted from 4×4 submatrices of the saddle matrix.

In order to detect variations in compartments between groups, the replicate PC1 values for each age group were used as an input for lmFit/eBayes functions from the *limma* v.3.62.2^33^ R package. This analysis aimed to identify regions demonstrating significant alterations in PC1 values (with an FDR < 5% and a minimum absolute change of 0.3 in PC1 value). GO annotation of identified differential bins was made using *clusterProfiler* R package v.4.14.6^34^, with all RefSeq genes as a background. Additionally, differential bins in A and B compartments were classified as “strong” (s) and “weak” (w) based on the relationships between their E1 values. A compartment bins with E1_target_ < E1_adult_ were classified as wA→sA transitions, while those with E1_target_ > E1_adult_ were classified as sA→wA transitions. The same stratification approach was applied to B-compartment bins, accounting for the sign of E1 values.

### Identification of topologically associating domains

TAD boundaries were determined at 10-kb resolution via *cooltools* using the insulation score method, with window size 50 kb. A TAD boundary was considered differential (unique) compared to wild type (WT) if its coordinates were altered from coordinates of WT by more than one bin (10 kb). The average TAD plot was generated via *Coolpup.py* package v.1.1.0^35^. Interaction frequencies of TADs were computed using the HiCExplorer package through hicAverageRegions function v.3.7.2^36^ (PMID: 29901812) with --range = 200000, -cb = ‘center’. Intensity profiles of TAD borders were analyzed by computing log₂-transformed contact intensity values which were extracted along the central TAD diagonal, spanning from the base to the apex of the triangular interaction matrix. Boundary strength was calculated based on the peak prominence algorithm using cooltools. Genes located near cell-specific TAD boundaries were determined using *ChIPseeker* R package v.1.42.1^37^ and GO annotation made using *clusterProfiler* R package v.4.14.6^34^, with all genes in TAD boundaries as a background.

### Chromatin loop calling

Loops were detected in 10-kb resolution via *Chromosight* v.1.6.3^38^ with parameters as follows: minimal loop size = 35kb, maximal loop size = 3.5Mb, Pearson correlation coefficient threshold = 0.45. Differential loops were identified as those falling into the top or bottom 10th percentile in terms of interaction intensity difference between two conditions and average loop profiles were generated via *GENOVA* R package v.0.95^39^. Loop strength for each condition was quantified according to the method proposed in Gassler et al.^40^ Briefly, we first calculated the average signal in the central 7×7 submatrix of the aggregated loop matrix and then subtracted from it the average signal from the combined upper-left and bottom-right submatrices of the same sizes. Differential loop strength (DLS) was calculated as the difference between loop strengths between two conditions. To assess statistical significance of this difference, we performed a permutation test by randomly shuffling condition labels of submatrix values, recalculating the loop strength difference for each permutation, and repeating this process 10000 times. The p-value was calculated as the number of cases with label shuffling where the strength difference is as extreme as or more extreme than the observed value (plus one), divided by the total number of permutations.

Loops were annotated using *ChIPseeker* R package v.1.42.1^37^. For the analysis of distribution of regulatory elements within loop anchors FANTOM5 mm10 enhancer coordinates were utilized (https://fantom.gsc.riken.jp). Genomic loops were categorized based on differential contact frequencies relative to wild-type (WT): the top decile of loops with the greatest positive fold change were classified as up-folded, while the bottom decile (most negative fold change) were designated as down-folded. Enrichment analysis of histone marks or transcription factors within loop anchors was performed using neural ChIP-seq datasets from the ChIP-Atlas database^41^.

### RNA library preparation and sequencing

RNA was extracted from primary cortical cultures five days after infection with shSIRT6 / shCtrl, using Total RNA Purification Kit (NORGEN, cat.35350) according to the manufacturer recommendations. RNA sample QC, library preparations, sequencing reactions, and initial bioinformatic analysis were conducted at GENEWIZ, LLC./Azenta US, Inc (South Plainfield, NJ, USA). Total RNA samples were quantified using Qubit 4.0 Fluorometer (Life Technologies, Carlsbad, CA, USA) and RNA integrity was checked with 4200 TapeStation (Agilent Technologies, Palo Alto, CA, USA). Samples were initially treated with TURBO DNase (Thermo Fisher Scientific, Waltham, MA, USA) to remove DNA contaminants. ERCC RNA Spike-In Mix (Cat: #4456740) from ThermoFisher Scientific, was added to normalized total RNA prior to library preparation following manufacturer’s protocol. The next steps included performing rRNA depletion using QIAGEN FastSelect rRNA HMR or Fly Kit (Qiagen, Germantown, MD, USA), which was conducted following the manufacturer’s protocol. RNA sequencing libraries were constructed with the NEBNext Ultra II RNA Library Preparation Kit for Illumina by following the manufacturer’s recommendations. Briefly, enriched RNAs are fragmented for 15 minutes at 94°C. First strand and second strand cDNA are subsequently synthesized. cDNA fragments are end repaired and adenylated at 3’ends, and universal adapters are ligated to cDNA fragments, followed by index addition and library enrichment with limited cycle PCR. Sequencing libraries were validated using the Agilent Tapestation 4200 (Agilent Technologies, Palo Alto, CA, USA), and quantified using Qubit 4.0 Fluorometer (ThermoFisher Scientific, Waltham, MA, USA) as well as by quantitative PCR (KAPA Biosystems, Wilmington, MA, USA). The sequencing libraries were multiplexed and clustered on the flowcell. After clustering, the flowcell was loaded on the Illumina NovaSeq instrument according to the manufacturer’s instructions. The samples were sequenced using a 2×150 Pair-End (PE) configuration.

### RNA-seq data processing and analysis

RNA-seq data from generated shSirt6 and shCtrl samples, as well as public CK-p25 dataset of Step 1, Step2 and Controls replicates^42^ was processed using the nf-core/rnaseq pipeline v3.16.0^43^. Obtained .fastq files were first trimmed with Trim Galore! and Cutadapt and then aligned to the mm10 reference genome with STAR. Transcript expression was quantified with Salmon and next summarized to gene-level counts via *tximport* package v1.34.0^44^. In the shSIRT6/shCtrl dataset one shSIRT6 replicate was excluded due to poor quality control metrics. Differential expression analysis was conducted with DESeq2 v1.46.0^45^, applying significance thresholds of FDR p-value < 0.05. Jaccard similarity between up- and down-regulated DEG sets from each comparison were calculated using GeneOverlap R package v1.42.0^46^. For functional annotation, protein-coding genes meeting these criteria were subjected to Reactome pathway enrichment analysis using the enrichPathway function from ReactomePA v1.50.0^47^ with the following parameters: pvalueCutoff = 0.05.

Publicly available snRNA-seq data from Old and juvenile Control mouse brains^48^ was used to assess age-related transcriptional alterations in mouse neurons. First, we converted already processed and filtered cell-specific expression profiles to pseudobulk count matrix via ADPBulk Python package v0.1.4^49^. The resulting matrix was filtered to retain only “inhibitory interneuron”, “medium spiny neuron” and “neuron” cell types, selecting donors of 4 week old and 90 week old. The remaining analyses were performed using the same methodology as described above.

### Immunofluorescence

Cells were rinsed with phosphate buffer saline (PBS) and fixed with 4% paraformaldehyde for 10 min at room temperature, followed by two additional washes. Cells were permeabilized (0.1% tri-sodium citrate and 0.1% Triton X-100 in Distilled water, pH 6) for 5 min and rinsed again. After 30 min of blocking (0.5% bovine serum albumin [BSA], 5% goat serum, 0.1% Tween-20 in PBS), cells were incubated with SIRT6 antibody (abcam, Ab88494) diluted 1:200 in blocking buffer overnight at 4°C. The next day, cells were washed three times with wash buffer (0.25% BSA, 0.1% Tween-20 in PBS), incubated for 1h with the secondary antibody (goat anti rabbit Alexa Fluor Plus 555, Thermo Fisher Scientific, A32732, diluted in blocking buffer 1:200) at room temperature and rinsed three more times. Cells were then DAPI-stained for three minutes at room temperature and rinsed with PBS twice before imaging. Nuclear mean intensity was measured and analyzed using ImageJ (https://imagej.net/ij/download.html).

### Nuclear morphology analysis

Nuclei were isolated from young (4-5 months), middle-aged (10-13 months), and old (18.5-20 months) S6-KO mice and processed following the same staining protocol as their WT littermates, as described in Kriukov et al^10^. During image analysis, only NeuN-positive nuclei were selected, and nuclear area was quantified using CellProfiler^50^ with the same settings described in the reference above. To assess the statistical significance of nuclear area differences between WT and S6-KO mice across ages, a linear mixed-effects model was fitted: *Area ∼ Genotype × Age + (1|Mouse_ID)*, followed by a two-way ANOVA.

### Analysis of TF-target interactions

The BART3D tool v.1.1^51^ was used to identify transcription factors (TFs) associated with differential chromatin interactions (DCIs). Transcription factors associated with differentially expressed genes (DEGs) were identified using enrichment analysis via the enrichR package v.3.4^52,53,54^. DEGs were cross-referenced against the ChEA3 database^55,56^, of experimentally validated TF-target interactions, with significance defined as an FDR p-value < 0.05.

## Results

### SIRT6 deficiency promotes chromatin decompaction through a mechanism independent of aging

To assess alterations in chromatin folding upon SIRT6 deficiency, we generated Hi-C maps for mouse neuronal nuclei isolated using FANS from the cortex of brain-specific SIRT6-knockout (S6-KO) and wild-type (WT) mice^28^. We analyzed adult S6-KO mice (10-11 months, N=3), adult WT mice (10 months, N=3), and old WT mice (20 months, N=3) (Fig.1a, Supplementary Table 1). The principal component analysis performed on compartment scores reveals distinct segregation between adult WT, old WT, and adult S6-KO neurons (24% of the explained variance, Fig.1b), suggesting significant differences in chromatin organization between adult WT, old WT, and adult S6-KO neurons. S6-KO-associated changes are predominantly captured along PC1, while aging-associated changes (old vs. adult) are separated along PC2. The orthogonality of these principal components indicates that SIRT6 loss induces chromatin restructuring distinct from aging, suggesting a unique mechanism rather than physiological aging.

The analysis of interactions within all chromosomes (cis-contacts) and between all pairs of chromosomes (trans-contacts) performed for Hi-C maps demonstrates a significant decrease in trans-contacts and a significant increase in cis-contacts in both old WT and adult S6-KO neurons compared to adult WT neurons (Wilcoxon rank sum test p-value < 10^-5^ in both cases, Fig.1c). These alterations indicate similar patterns of chromosome territory expansion in both S6-KO and old WT cells compared to adult WT cells. In line with our prior work, in which we demonstrated an expansion in cell nuclei size in old WT neurons^10^, we measured S6-KO mice nuclei, and also demonstrate age-related enlargement of nuclei compared to WT neurons (Fig.1d, Supplementary Fig. 1a,b).

In order to reveal the properties of the chromosome fiber under SIRT6 deficiency, we plotted the average interaction frequency over genomic distance (Fig.1e). In all three groups (adult WT, old WT and S6-KO), the autosomal chromatin contact probability *P(s)* with genomic distance *s* exhibits a power-law scaling with the slope of *∼s^−1^*, consistent with the chromosome folding mode of fractal globules^57,58^. The *P(s)* ratio plot calculated between old and adult WT cells shows that chromatin in aging neurons is more compact at local linear distances and less compact at long-range distances. Conversely, the *P(s)* ratio plot calculated between S6-KO and adult WT neurons demonstrates a slight decrease in short-range chromatin interactions and a prominent increase in long-range interactions upon S6-KO.

To estimate the significance of these differences and to evaluate chromatin compaction in the models, we employed two metrics: the ratio of distal (>10.5 Mb) to local (<10.5 Mb) Hi-C cis-interactions (referred to as DLR), and the proportion of trans-interactions, relative to the overall number of interactions observed at a genomic locus (referred to as ICF). A notable decrease in DLR in aging neurons highlights a significant transition of chromatin interactions towards shorter distances in older neurons (Wilcoxon test p-value < 10^-5^, Fig.1f). This result is consistent with the observed increase in the contact frequency ratio at short-range genomic distances for old WT neurons (Fig. 1e). Similarly, a significant increase in DLR upon S6-KO (Wilcoxon rank sum test p-value < 10^-5^, Fig.1f) supports the increased contact frequency ratio at long-range genomic distances for S6-KO neurons (Fig. 1e), indicating a transition towards long-range chromatin interactions. Interestingly, a significant decrease in ICF (Wilcoxon test p-value < 10^-5^, Fig. 1g) is observed in both aged and S6-KO neurons compared to adult WT (average ΔICF = 0.86 for old WT vs. adult WT and 0.96 for S6-KO vs. adult WT). This ICF reduction suggests that chromatin decompaction in aging and SIRT6 deficiency preferentially facilitates intrachromosomal interactions while decreasing interchromosomal contact frequency, as supported by cis/trans comparisons (Fig. 1c).

**Figure 1.**
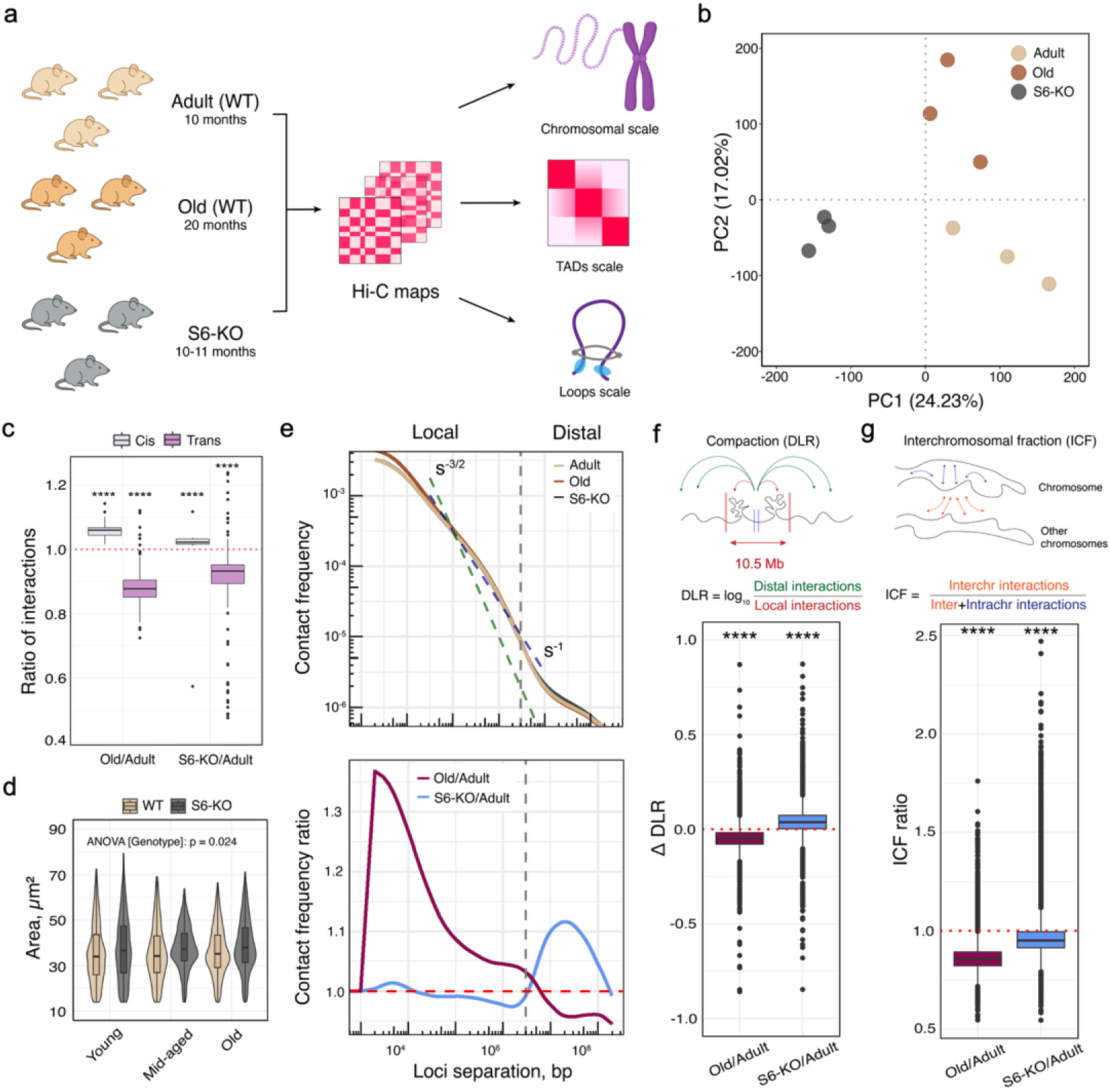
Genome-scale changes in chromatin folding in aging and S6-KO mouse neurons. a. Study design. Hi-C contact matrices were generated from neurons of old WT, adult WT, and adult S6-KO mice, followed by multilevel analysis of chromatin organization. b. PCA plot based on eigenvector (E1) values. Colors represent adult S6-KO, adult WT and old WT replicates. c. Average ratios of cis-interactions (gray) and trans-interactions (pink) calculated for old WT neurons relative to adult WT (left) and for adult S6-KO neurons relative to adult WT (right). d. Nuclear area measurements for young, middle-age and old WT and S6-KO mice. e. Upper panel: Polymer scaling P(s) curves showing average interaction frequencies across genomic distances s. Dashed lines represent the slope of a fractal globule (∼ s^−1^) and the slope of an equilibrium globule (∼ s^-3/2^). Lower panel: Scaling ratio plot showing the ratio of average interaction frequencies across genomic distances calculated for old WT relative to adult WT neurons (brown line) and for S6-KO relative to adult WT neurons (blue line). f. Differences in distal (>10.5 Mb) to local (<10.5 Mb) contact ratios (DLRs) calculated between old and adult WT neurons, and between S6-KO and adult WT neurons. g. ICF ratios calculated for old WT relative to adult WT neurons, and for S6-KO relative to adult WT neurons. In all panels, asterisks indicate Wilcoxon rank sum test p-value < 0.00001.

Overall, our results demonstrate that S6-KO neurons, despite sharing reduced interchromosomal interactions with aged neurons, are characterized by generally distinct chromatin folding patterns linked to overall chromatin decompaction.

### SIRT6-deficient neurons demonstrate differential chromatin compartment patterns

Long-range contact frequencies vary between old and adult WT neurons, as well as between S6-KO and adult WT neurons, suggesting potential differences in chromatin compartmentalization across these conditions. Notably, both visual inspection and eigendecomposition of Hi-C maps (Fig. 2a) indicate distinct positioning and compartmentalization of chromatin in aging and S6-KO neurons compared with adult WT cells. In both old and S6-KO mice, the B compartment shows increased interaction strength, while the A compartment shows reduced strength. This suggests that inactive chromatin (B) gains more interactions in old and S6-KO neurons, whereas active chromatin (A) loses some interactions.

**Figure 2:**
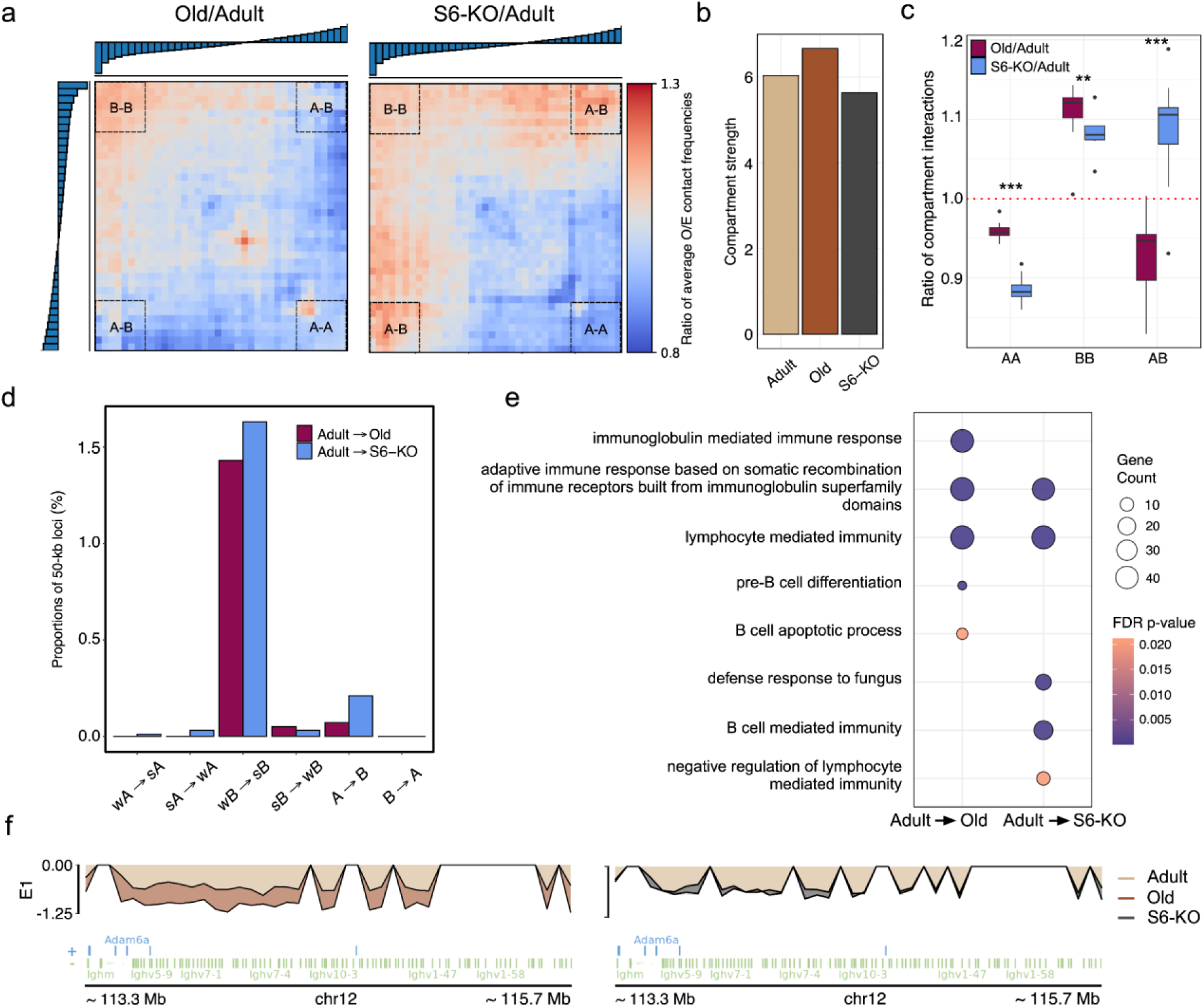
Compartmentalization changes in aging and S6-KO mouse neurons. a. Ratios of saddle plots (observed-over-expected contact frequencies of genomic regions sorted by their E1 rank) between conditions: old/adult WT (left) and S6-KO/adult WT (right). b. Compartment strength, calculated as the ratio of intra-compartment to inter-compartment interactions. c. Ratios of A-A, B-B and A-B compartment interactions calculated between conditions: old/adult WT (purple) and S6-KO/adult WT (blue). d. Proportions of 50-kb loci across the genome showing various compartment transitions in adult WT → old WT and adult WT → S6-KO comparisons. “Strong” (s) and “weak” (w) indicate the magnitude of E1 values. e. GO enrichment analysis of genes overlapping wB → sB transitions. f. E1 profiles for differential 50-kb loci overlapping the immunoglobulin heavy chain variable region (*Ighv*) gene cluster on chromosome 12 in old WT and S6-KO neurons compared to adult WT.

Interestingly, the direction of changes in A–B compartment interactions is opposite between old and S6-KO mice (Fig. 2a-c). In old mice, there is a decrease in A-B interactions, suggesting stronger segregation between A and B compartments (Fig. 2b). In contrast, S6-KO mice demonstrate an increase in A-B interactions, which indicates weaker compartmentalization in S6-KO neurons (Fig. 2b,c).

In agreement with previous reports, compartments do not undergo global changes during either aging or S6-KO^10^. Specifically, analysis of the first eigenvector sign and amplitude reveals that only 0.07% of 50-kb loci in aging and 0.21% under S6-KO flip between active (A) and inactive (B) compartments (A→B or B→A). These loci were identified based on statistically significant changes in eigenvector values (FDR < 5%, |ΔE1| > 0.3).

We next explored changes in the magnitude of the first eigenvectors between groups and observed that 1.53% of 50-kb loci in aging and 1.76% under S6-KO demonstrate significant shifts within the same compartment type (within A or B; FDR < 0.05 and |Fold Change| > 0.3). In each group, 50-kb loci were further classified as “strong” (sA or sB) or “weak” (wA or wB) based on E1 values (see Methods). In both comparisons (adult WT→old WT and adult WT→adult S6-KO), wB→sB and A→B transitions prevail. Specifically, 1.48% of loci in aging and 1.69% under S6-KO exhibit wB→sB transitions, compared to 0.05% and 0.03% sB→wB transitions, respectively (Fig. 2d). Similarly, 0.08% of loci in aging and 0.22% under S6-KO demonstrate A→B transitions, compared to 0% and 0.004% B→A transitions, respectively (Fig. 2d). These patterns suggest that both aged and S6-KO mouse neurons harbor a modestly larger fraction of their genome in the B compartment.

We next identified genes located within genomic regions demonstrating compartment transitions (wA→sA, sA→wA, wB→sB, sB→wB, A→B, and B→A). Functional enrichment analysis of genes from wB→sB compartment transitions in both SIRT6 deficiency and aging revealed immune-related terms, including adaptive immune response, lymphocyte mediated immunity, and B-cell mediated signaling (Fig. 2e, Supplementary Tables 3,4). Notably, these terms were driven by a cluster of genes on chromosome 12 encoding immunoglobulin heavy chain variable region (*Ighv*) subunits (59 genes in aging neurons, 38 genes in S6-KO neurons). This genomic region undergoes wB → sB transition with a pronounced decrease in the first eigenvector values in aging and S6-KO compared to adult WT (Fig. 2f). Thus, there is a remarkable wB → sB compartment transition, particularly evident in B compartmentalization increase of the genome region harboring immunoglobulin heavy chain variable region (*Ighv*) genes located on chromosome 12. In neurons, the *Ighv* locus remains transcriptionally silent through repressive histone modifications and CTCF/cohesin-mediated insulation blocking VDJ recombination on both alleles^59^. Given the reduced activity of SIRT6 and CTCF in aging and AD^1,60^, the observed strengthening of B compartmentalization at Ighv locus could be reflected by potentially weakened chromatin insulation and consequent compensatory heterochromatinization.

Taken together, we found that chromatin in old WT and S6-KO mouse neurons differs in the extent of segregation between A and B compartments, but exhibits similar changes in compartmentalization within A and B compartments compared to adult WT, most of which are driven by wB→sB transitions, predominantly involving immune response genes.

### SIRT6 deficiency leads to less pronounced TADs in mouse cortical neurons

To analyze changes in TAD organization upon SIRT6 depletion and aging, we annotated TADs using the IS algorithm^31^ and identified 6230 TAD boundaries shared among adult WT, old WT, and S6-KO neurons (Fig. 3a). In addition, 2132 (19.3%) boundaries were specific to old WT neurons, 1707 (15.0%) to S6-KO neurons, and 1768 (15.4%) to adult WT neurons. On average, TADs were more pronounced in old WT but less prominent in S6-KO neurons compared to adult WT (Fig. 3b-d). The stronger contact intensity at the TAD level in aged neurons corresponds to the observed overall enrichment of short-range chromatin interactions, whereas the reduced contact intensity in S6-KO neurons reflects a depletion of these interactions (Fig. 1d).

**Figure 3:**
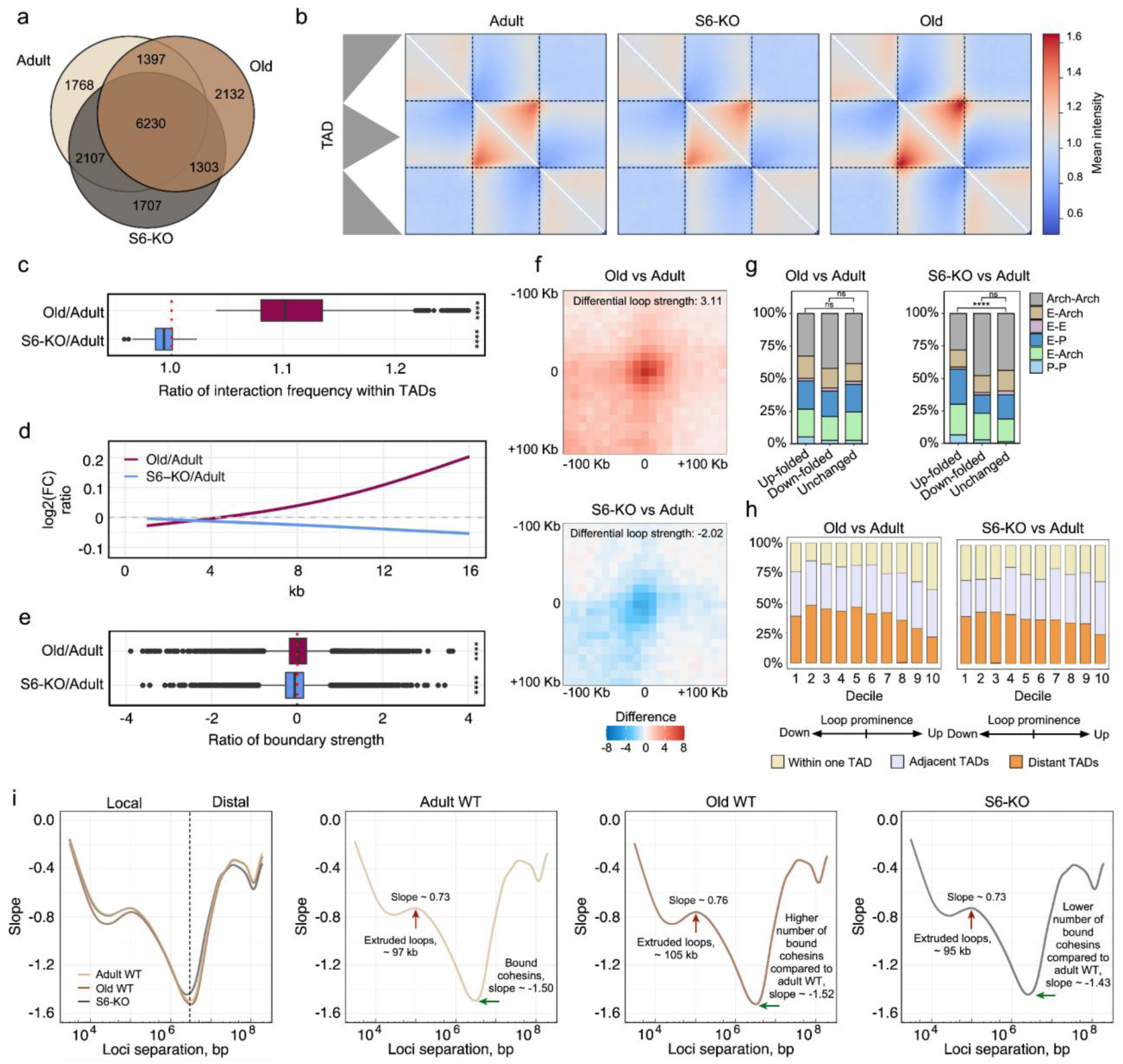
Differences in TAD organization and loop formation in aging and S6-KO neurons. a. Venn diagram showing the overlap of TAD boundaries in adult WT, old WT and S6-KO neurons. b. Average TAD plots for adult WT, S6-KO, and old WT. c. Ratios of interaction frequencies within TADs for old/adult WT (purple) and S6-KO/adult WT (blue) neurons. Asterisks indicate Wilcoxon Rank Sum Test p-values: **** - p<0.0001. d. Log_2_ ratios of interaction intensities within TADs for old/adult WT and S6-KO/adult WT comparisons, computed at different genomic distances separately to illustrate distinct scaling patterns. A schematic explaining the methodology of calculation is shown in Supplementary Fig. 4. e. Ratio of strength for boundaries common between Old and Adult WTs, S6-KO and Adult WT. Asterisks indicate Wilcoxon Rank Sum Test p-values: **** - p<0.0001. f. Average loop intensity differences between old WT and adult WT neurons (left) and between S6-KO and adult WT neurons (right). The color scale shows the difference in the number of contacts between average loops calculated for each condition. g. Up-folded, down-folded and unchanged loops in aging (left) and S6-KO (right), classified by the presence of regulatory elements (promoters or enhancers) within loop anchors or their absence (architectural loops). Asterisks indicate χ² test p-values for differences in loop-class proportions: **** - p < 0.0001 h. Loops stratified into deciles based on magnitude and direction of change in aging (left) and S6-KO (right), classified by their anchor positions: within a single TAD (yellow), spanning two adjacent TADs (purple), or bridging two distant TADs (orange). i. Slopes of log–log Pc(s) curves for each condition. Vertical red arrows mark the maximum slope position (mean cohesin-extruded loop size), and horizontal green arrows indicate the minimum slope depth (density of cohesin occupancy).

Additionally, old WT neurons show a slight increase in mean and median TAD size compared to adult WT, suggesting some degree of TAD expansion (Supplementary Fig. 2). At the same time, the strength of TAD boundaries is lower in S6-KO cells, indicating weakened TAD segregation (Fig. 3e). Although TAD sizes in S6-KO neurons are comparable to those in aged cells (Supplementary Fig. 2), their overall TAD prominence is reduced, potentially due to a global decrease in short-range chromatin interactions.

We next examined the functional relevance of genes located at condition-specific TAD boundaries. GO enrichment analysis revealed that TAD boundaries unique to old WT neurons are significantly enriched for genes involved in cilium organization (FDR p-value = 1.1227 × 10^-3^) and neuron migration (FDR p-value = 1.1227 × 10^-3^), while S6-KO-specific TAD boundaries are significantly enriched for genes regulating synapse assembly (FDR p-value = 1.2126 × 10^-5^) (Supplementary Fig. 3, Supplementary Tables 5,6), which also indicates that aging and SIRT6 depletion reshape TAD organization in different ways.

### Lack of SIRT6 induces differential chromatin looping in mouse cortical neurons

To investigate regulatory interaction changes under SIRT6 depletion or aging, we further analyzed differential chromatin looping. In total, 4489, 5050 and 5949 Hi-C loops were annotated in S6-KO, adult WT and old WT neurons, respectively. Overall, old WT neurons exhibit more pronounced loops (differential strength = 3.11, permutation test p-value < 1 × 10^-4^) compared with adult WT, whereas S6-KO samples show less pronounced loops (differential strength = −2.12, permutation test p-value < 1 × 10^-4^) (Fig.3f). These findings are consistent with the observed enrichment of short-range chromatin interactions (<2 Mb) in aged cells and depletion of these interactions in S6-KO neurons (Fig. 1d).

Out of the union of 11054 loops annotated in S6-KO, adult WT or old WT neurons, the top 10% of loops showing the largest intensity increase or decrease in S6-KO compared to adult WT were classified as up- or down-folded, respectively. Up- and down-folded loops in old WT compared to adult WT were defined similarly. Loops down-folded under SIRT6 deficiency show significant enrichment for histone marks associated with enhancers and active genes – H3K27ac (q-value = 6.4×10⁻¹²) and H3K4me3 (q-value = 2.6×10⁻⁷). Notably, the same marks are also enriched in the anchors of loops down-folded in aged cells (q-value = 8.29×10^⁻¹²^ and 2.46×10⁻⁶, respectively), indicating that SIRT6 loss recapitulates age-related loop remodeling (Supplementary Tables 7,8). Functional analysis of genes located within anchors of these loops did not reveal specific pathway enrichment, suggesting a general decline in enhancer-mediated transcriptional control as a shared feature of both SIRT6 deficiency and aging.

We further analyzed the distribution of regulatory elements (enhancers and promoters) within the anchors of up- and down-folded loops. A substantial fraction of down-folded loops in both aging and S6-KO lacks regulatory elements at their anchors and appears to serve primarily architectural functions. Furthermore, in both aged and S6-KO neurons, the proportion of up-folded loops containing regulatory elements increased compared to down-folded and unchanged loops. Moreover, in SIRT6-KO, significant differences were observed in the distribution of loop types relative to unchanged loops (p-value = 2.47 × 10⁻⁸, χ² test), primarily driven by an increased fraction of enhancer–promoter interactions in S6-KO. (Fig. 3g).

We next examined the spatial distribution of up- and down-folded loops by classifying them according to the position of their anchors relative to TADs: loops confined within a single TAD, spanning two adjacent TADs, or bridging two distant TADs. In both aged and S6-KO neurons, up-folded loops are predominantly confined within a single TAD or span adjacent TADs, whereas down-folded loops more often connect distant TADs (Fig. 3h). Taken together, these findings suggest that, in both aging and S6-KO, regulatory loops in the same or adjacent TADs tend to be unchanged or even up-folded, while architectural loops spanning distant TADs are disrupted.

We further characterized differential loop formation in S6-KO, adult and old WT neurons by analyzing the average size of chromatin loops. Previous work has shown that quantitative derivative analysis of log-log Pc(s) curves identifies the mean cohesin-extruded loop length at the inflection point (maximum derivative/minimum slope), while the subsequent local minimum depth reflects cohesin linear density^40^. We performed this analysis and observed larger mean loop sizes in aged neurons compared with adult (∼105 kb vs. ∼97 kb) and smaller mean loop sizes in S6-KO neurons (∼95 kb vs. ∼97 kb). Consistently, these shifts were accompanied by increased cohesin occupancy in old relative to adult neurons and reduced cohesin binding in S6-KO (Fig. 3i).

Taken together, our results demonstrate that SIRT6 deficiency disrupts chromatin architecture by weakening loop strength, shortening loops, and promoting potentially aberrant long-range interactions between distant TADs. In contrast, aging promotes the chromatin loop formation associated with increased cohesin occupancy in mouse neurons.

### Brain SIRT6 Deficiency and AD-like models alter chromatin structure differently from normal aging

SIRT6 levels decline in the brain during physiological aging^23,24^ and are also reduced in postmortem brain samples from individuals with Alzheimer’s disease (AD)^1,2,3,4,25,61,62^. Multiple studies suggest a protective role of SIRT6 against AD pathology^2,3,61^. To explore the 3D genome regulatory consequences of SIRT6 reduction in physiological and pathological aging, we compared chromatin architecture in the aging brain and S6-KO neurons with AD-like conditions. Specifically, we analyzed Hi-C data derived from CK-p25 mice, a well-established AD model^63^. We chose this model since in this study neurons were classified according to levels of the neuronal nuclei marker NeuN and DNA damage marker γH2AX into three groups: *Control neurons* (high NeuN, baseline γH2AX), *Step 1 neurons* (high NeuN, high γH2AX), and *Step 2 neurons* (low NeuN, high γH2AX)^19^, an architectural deterioration increased with damage burden, supporting an association between DNA damage and 3D genome breakdown. While increased DNA damage is well documented in SIRT6 deficient brains, our Hi-C study did not quantify DNA damage, therefore the damage-architecture relationship is currently correlative across models.

Our analysis revealed a notable decrease in interactions between A and B chromatin compartments in both Step 1 and Step 2 neurons, similar to old WT neurons (Fig. 4a, b). This reduced intermingling suggests that enhanced segregation of A and B compartments is a common feature of aging and progressive neurodegeneration. Consistently, compartment strength analysis demonstrated progressively stronger compartmentalization in Step 1 and Step 2 neurons compared to controls, with fold changes of 1.13 and 1.15, respectively, versus 0.93 in S6-KO and 1.10 old WT neurons relative to adult WT (Figure 4a).

**Figure 4:**
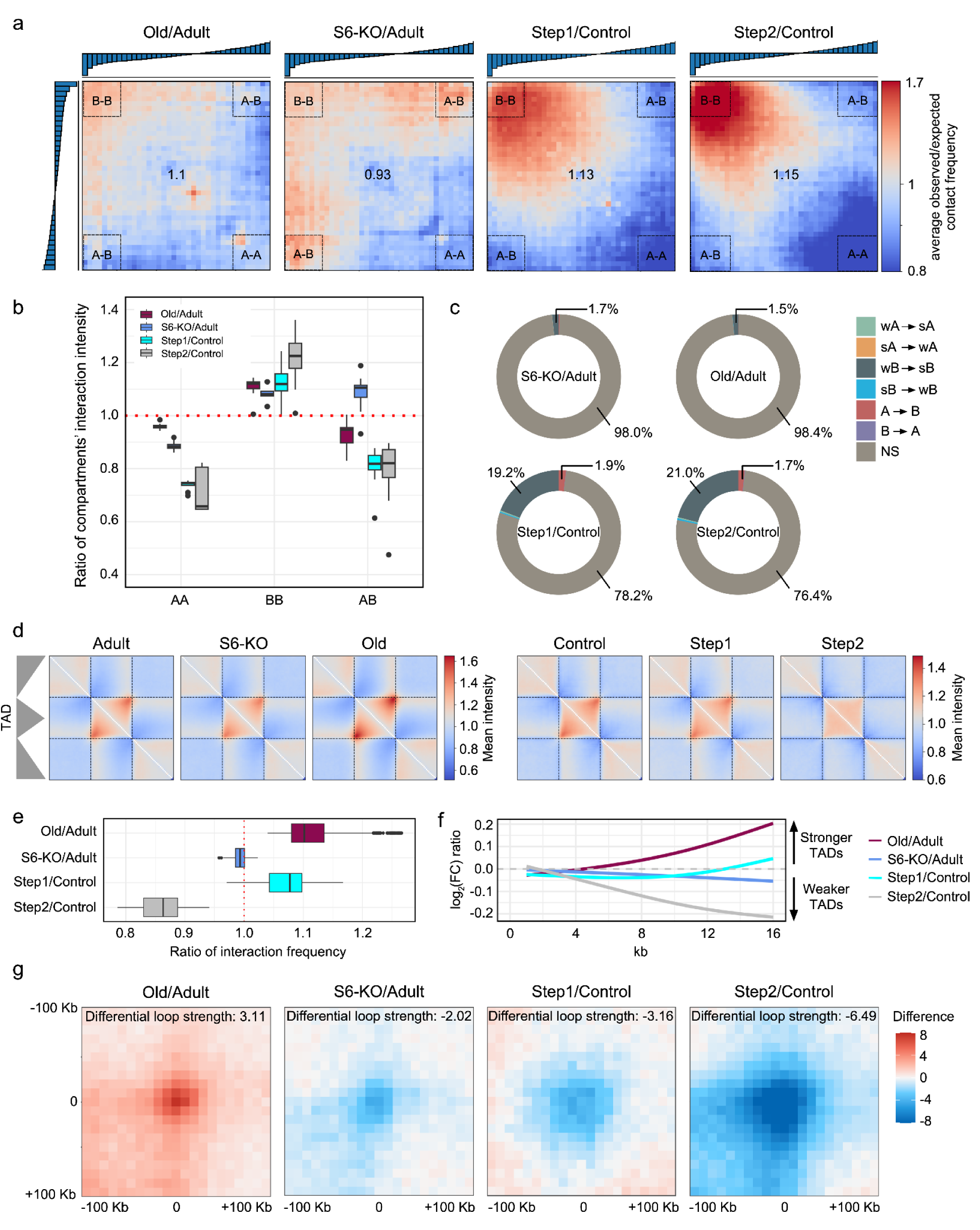
Chromatin architecture changes in SIRT6 deficiency, aging, and neurodegeneration. a. Saddle plots showing ratios of observed/expected contact frequencies (Old/Adult WT, S6-KO/Adult WT, Step 1/Control and Step 2/Control), arranged by E1 rank. Values in the center of each plot represent compartment strength. b. Ratios of A-A, B-B and A-B interactions across the same comparisons. c. Donut charts illustrating proportions of differential 50-kb bins compartment transitions (Adult → Old, Adult → S6-KO, Control → Step 1, Control → Step 2). “Strong” (s) and “weak” (w) indicate the magnitude of the first eigenvectors. d. Average TAD plots across experimental conditions. e. Boxplots of ratios of interaction frequencies within TADs across the same comparisons. f. Log_2_ ratios of contact intensities at TAD borders across the same comparisons, calculated from the center of the bottom to the top of the central TAD triangle (explained in Supplementary Fig. 4), highlighting distinct scaling patterns. g. Ratios of average loop plots across the same comparisons. The color scale shows the difference in the number of contacts between average loops calculated for each condition.

To further explore compartmental remodeling in neurodegeneration, we analyzed compartmental shifts in Step 1 and Step 2 neurons. Specifically, 22% and 24% of 50-kb loci, respectively, demonstrated shifts between A and B compartments or changes within the same compartment type (weak to strong or strong to weak). The most prevalent transitions were from weak B to strong B (wB→sB), affecting 19.21% of loci in Step 1 and 21.03% in Step 2, followed by A→B transitions (1.89% and 1.68%, respectively) (Fig. 4c). This pattern mirrors our observations in aged and S6-KO neurons, indicating that expansion of the B compartment represents a shared hallmark of chromatin remodeling during both aging and neurodegeneration (Supplementary Table 9).

To understand the functional consequences of these compartmental shifts, we performed enrichment analysis of genes associated with wB→sB transitions. This analysis revealed significant enrichment of terms related to adaptive humoral immunity, including pre-B-cell activation, humoral immune response, B-cell activation, and B-cell signaling (Supplementary Fig. 4, Supplementary Table 10). A prominent cluster on chromosome 12, encompassing genes encoding subunits of the immunoglobulin heavy chain variable region (*Ighv*), was consistently enriched in these transitions in Step1 neurons, mirroring the patterns observed in S6-KO and old WT mice. In Step 1 neurons, 49 *Ighv* genes were involved, with similar numbers identified in old WT and S6-KO neurons (N = 59 and 38, respectively; Supplementary Fig. 6, Supplementary Table 11). Together, these results highlight a recurrent wB→sB transition linked to *Ighv* downregulation of *Ighv* genes across aging, SIRT6 deficiency, and neurodegeneration.

Furthermore, we compared TADs across S6-KO, aging, and neurodegeneration models. Density of contacts within TADs was increased in Step 1 neurons compared to Control neurons and in old WT compared to adult WT, but decreased in Step 2 and S6-KO neurons compared to their respective controls (Fig. 4d,e). Consistently, TAD boundary strength decreased in Step 2 and S6-KO cells but increased within average TAD corner regions in Step 1 neurons, similarly to old WT neurons (Fig. 4f). These findings suggest that early neurodegeneration and normal aging are associated with transiently more prominent TADs, whereas SIRT6 loss and advanced neurodegeneration lead to their erosion.

Similar to S6-KO neurons, both Step 1 and Step 2 neurons exhibited less pronounced loops compared to Control neurons (differential loop strength = −3.16 and 6.49, permutation p-value < 1 × 10^-4^, in Step 1 and Step 2 neurons, respectively) (Fig. 4g). Loop classification analysis revealed a consistent pattern of chromatin reorganization across SIRT6 deficiency and neurodegeneration. Architectural loops (not linked to regulatory elements) were predominantly lost (i.e., down-folded) in both Step 1 and Step 2, while the proportion of up-folded loops containing regulatory elements increased compared to down-folded and unchanged loops, with significant differences in loop type proportions detected in up-folded loops compared to unchanged ones (p-value = 2.47 × 10⁻⁸ in S6-KO, p-value = 9.18 × 10^-4^ in Step1, p-value = 4.32 × 10^-3^ in Step2, χ² test) (Supplementary Fig. 7). This suggests a shared shift in chromatin architecture across all conditions (aging, S6-KO, Step 1 and Step 2), where structural loops deteriorate as regulatory interactions tend to persist or even become more prominent.

### SIRT6 deficiency shows transcriptional alterations that overlap with those in AD-like models

To investigate transcriptomic alterations associated with chromatin structure remodeling in SIRT6-deficient neurons, we conducted an additional RNA sequencing experiment on cortical primary neurons from scrambled controls (WT, n=3) and shRNA-mediated SIRT6 knockdown (shSIRT6, n=2) mice. For comparison with normal and pathological aging, we also analyzed available gene expression datasets from CK-p25 AD model (Control, Step1, Step2; n=2 per group)^42^ and aging mice (Control, Old; n=2 per group)^48^. First, we verified SIRT6 silencing by immunofluorescence staining and observed a ∼50% reduction in SIRT6 protein levels (Supplementary Fig. 8). Next, we performed a PCA demonstrating clear separation between experimental and control samples, with the first principal component explaining 88% of the variance in Old vs. Control, 47% in shSIRT6 vs. shCtrl, and 66% in Step1/Step2 vs. Control (Fig. 5a).

**Figure 5:**
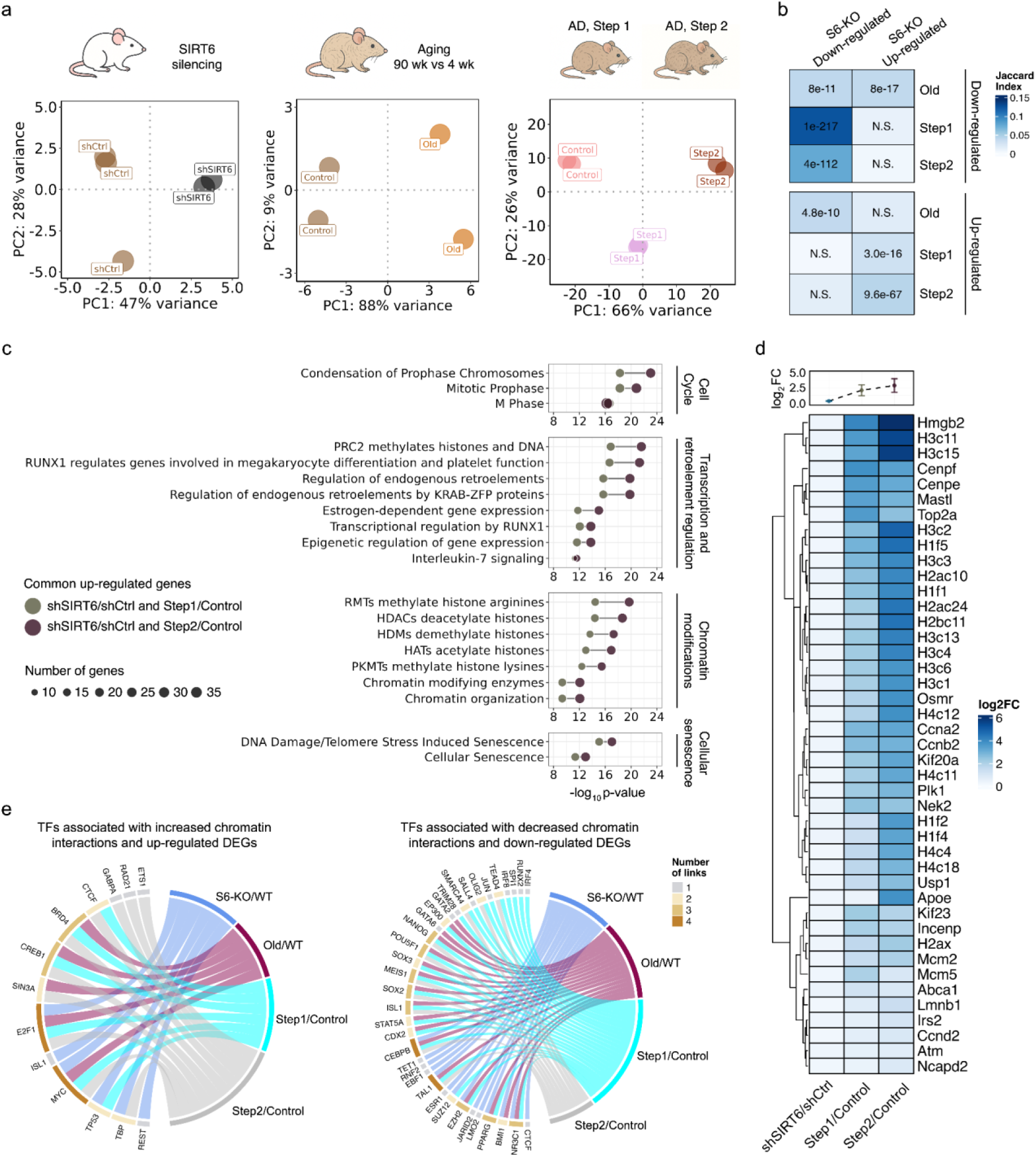
shSIRT6 neurons exhibit a transcriptional shift toward neurodegeneration. a. PCA plots showing separation between Control and Old, shCtrl and shSIRT6, Control and Step1/ Step2 transcriptomic profiles. b. Heatmap of Jaccard similarity between up- and downregulated DEGs across the same comparisons. Numbers indicate Fisher’s test p-values calculated for each intersection between gene sets. c. Dumbbell plot of top 20 significantly enriched Reactome terms among upregulated DEGs shared between shSIRT6/shCtrl and Step1/Control (olive green dots) or shared between shSIRT6/shCtrl and Step2/Control (dark purple dots). d. Heatmap of log_2_ fold changes for upregulated DEGs shared between shSIRT6/shCtrl, Step1/Control and Step2/Control. e. Circos plots showing overlap of TFs associated with upregulated genes and increased significant differential chromatin interactions in shSIRT6/shCtrl, Step1/Control and Step2/Control (left); overlap of TFs associated with downregulated genes and decreased differential chromatin interactions in S6-KO/WT, Old/WT, Step1/Control and Step2/Control (right). Color denotes degree of overlap: red – 2 groups; green – 3 groups; blue – 4 groups.

Differential expression analysis revealed strong overlap of downregulated DEGs between shSIRT6/shCtrl and both Step1/Control and Step2/Control (Fisher’s test p-values = 1.0 × 10^-217^ and 4.0 × 10^-112^, respectively), while overlap among upregulated DEGs was weaker but still significant (Fisher’s test p-values = 3.0 × 10^-16^ and 9.6 × 10^-67^, respectively) (Fig. 5b, Supplementary Tables 12-15). At the same time, shSIRT6/shCtrl and Old/Control mice exhibited similar DEG patterns but in opposite directions, consistent with the divergent alterations in chromatin organization observed in these two models.

To further explore functional similarities between SIRT6 deficiency and neurodegeneration, we performed Reactome enrichment analysis of DEGs shared between shSIRT6/shCtrl and Step1/Control or Step2/Control models. Shared upregulated genes were enriched for pathways related to chromatin post-translational modifications, regulation of transcription, cell cycle and cellular senescence, with enrichment generally stronger in shSIRT6 DEGs shared with Step 2 than with Step 1 (Fig. 5c, Supplementary Tables 16,17). Interestingly, genes in these pathways exhibited a progressive increase in expression from shSIRT6/shCtrl to Step1/Control and further to Step2/Control (Fig. 5d). A similar enrichment analysis of shared downregulated genes identified pathways related to neurotransmission functions and ion channel transport (Supplementary Tables 18,19), as well as a similar progressive trend of decreased expression of common genes in these pathways from shSIRT6/shCtrl to Step2/Control (Supplementary Fig. 9).

Next, we asked whether a common TF-mediated regulatory mechanism could explain the coordinated changes observed at both transcriptional and chromatin levels. To address this, we first used the BART3D method^51^ to identify TFs associated with significant changes in chromatin contact frequencies between experimental conditions. We next examined whether these TFs also regulate DEGs that demonstrate co-directional changes across the same conditions. Strikingly, REST emerged as a key TF associated with increased chromatin interactions and up-regulated DEGs in shSIRT6 neurons. REST is known as a repressor of neuronal gene transcription, with a remarkable role in neuronal synaptic plasticity, apoptosis, and neuroprotection in general. We previously demonstrated that SIRT6 facilitates interaction between REST and EZH2, a component of the Polycomb complex involved in epigenetic silencing and stem cell maintenance^64^, thereby derepressing REST targets^61^. Other enriched TFs included TBP, TP53, MYC, and E2F1 – known regulators of cell cycle progression, apoptosis, and stress responses^65,66,67,68^. They were consistently associated with increased chromatin interactions and upregulated DEGs in both SIRT6-deficient and neurodegenerative contexts. Furthermore, SUZ12, EZH2, and BMI1, also components of the Polycomb complex, were linked to upregulated interactions and DEGs in SIRT6 deficiency, aging, and neurodegeneration (Fig. 5f). Together, these findings suggest that SIRT6 deficiency promotes chromatin remodeling and transcriptional alterations through REST- and Polycomb-associated regulatory networks, reflecting changes observed in neurodegeneration.

Overall, in this study we observed both shared and divergent 3D changes that characterize aging and neurodegeneration in mouse models. All conditions exhibited some global chromatin alterations – a reduction in inter-chromosomal contacts (consistent with enlarged nuclei) and shifts towards a more inactive B-compartment state. However, key differences emerged: aged neurons maintained much of their local 3D genome organization (e.g. intact TAD structures and regulatory loops), whereas the S6-KO and CK-p25 neurons showed pronounced disruption of higher-order architecture (weakened TADs and loops). Transcriptomic analysis mirrored this pattern, with aging showing opposite changes compared to S6-KO and neurodegenerative models. Overall, these findings suggest that normal aging induces a “primed” chromatin state – displaying initial signs of reorganization – that may predispose neurons to further breakdown under pathological conditions, but on its own remains functionally compensatory.

## Discussion

Our study aimed to determine whether natural aging recapitulates early chromatin alterations observed in neurodegeneration or follows a distinct trajectory. By comparing cortical neurons from aged mice, S6-KO mice, and CK-p25 models, we found both shared and divergent changes in chromatin architecture and gene expression (Fig. 6).

**Figure 6:**
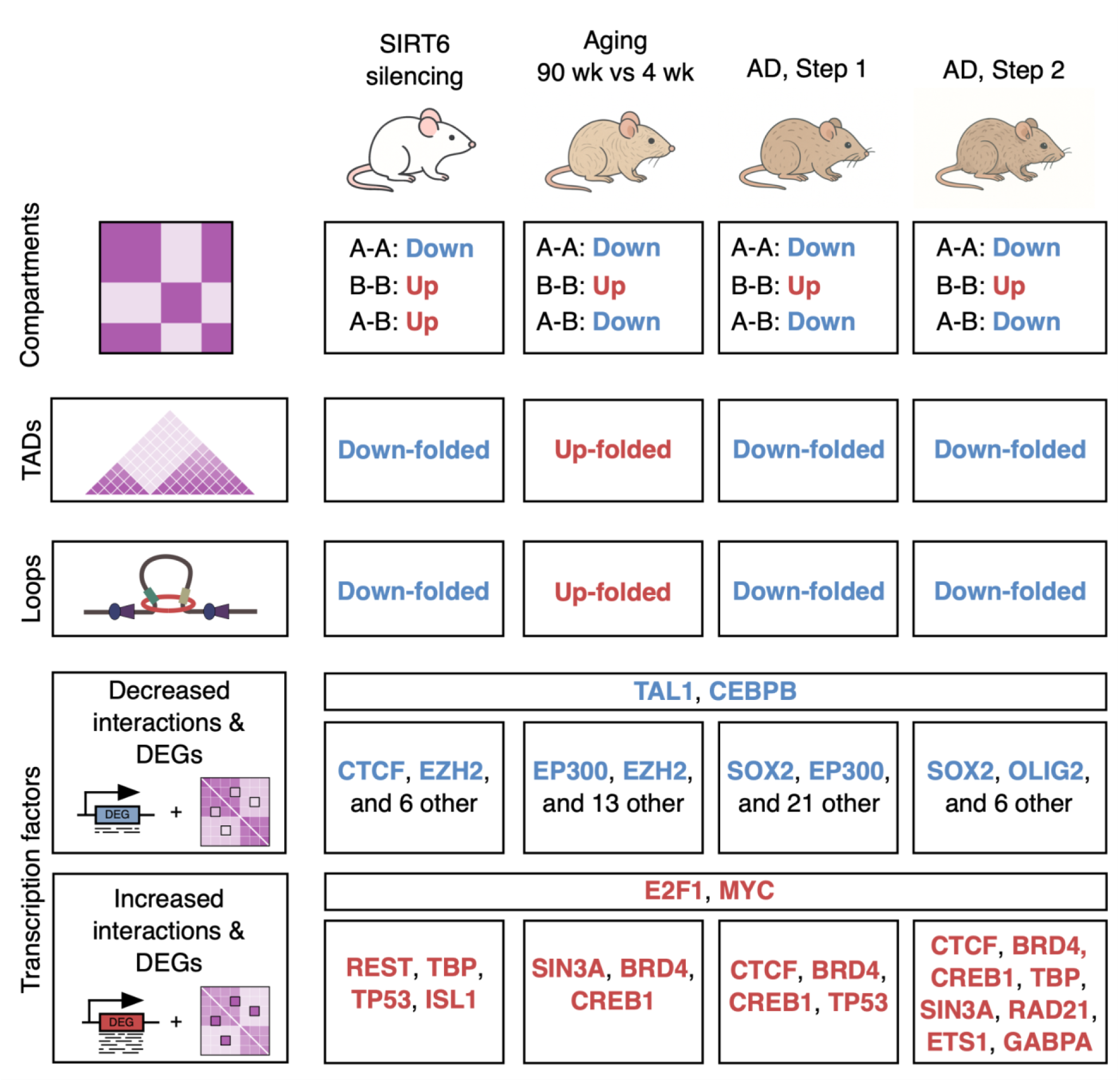
Summary of shared and divergent changes in chromatin architecture and gene expression in cortical neurons from aged mice, S6-KO mice, and CK-p25 models.

### Global chromatin architecture

Hi-C data demonstrated that adult, aged, and S6-KO cortical neurons form separate clusters, with S6-KO diverging orthogonally from the aging trajectory. Globally, both aged and S6-KO neurons exhibited fewer contacts between chromosomes, consistent with nuclear enlargement reported in both physiological neuronal aging and pathological states such as AD^10,69^. Aged neurons showed higher contact probabilities at short genomic ranges and fewer long-range contacts, whereas S6-KO neurons showed the opposite trend. Thus, normal aging increases local interactions (perhaps as a compensatory mechanism to preserve regulatory contacts), while SIRT6 loss causes a global disorganization that promotes aberrant long-range interactions. This analysis highlights that SIRT6 is critical for maintaining 3D genome integrity, and its loss induces architectural changes beyond those observed in physiological aging.

### Chromatin compartmentalization

A/B compartmentalization remained generally stable across all conditions, yet aged, S6-KO, and CK-p25 neurons exhibited strengthened B-B and weakened A-A interactions, indicating a shared trend towards a more repressive chromatin compartmentalization. But its prominence diverged, gradually increasing with the severity of the condition (S6-KO → Step 1 → Step 2). Importantly, the degree of A-B mixing also diverged: aging and neurodegeneration increased segregation of active and inactive domains, whereas S6-KO neurons showed greater A-B intermingling, implying loss of spatial compartment integrity.

Among affected loci, the immunoglobulin heavy chain variable region (Ighv) consistently shifted deeper into the B compartment in aged, S6-KO, and CK-p25 neurons. Normally, the Ighv locus is silenced in non-B cells by repressive histone modifications, DNA methylation, and CTCF/cohesin-mediated insulation^70,71,72^. CTCF and cohesin binding weakens with age^73^ and CTCF binding is diminished in AD cortex^60^. Together, these observations suggest that reduced activity of these architectural regulators may weaken chromatin insulation and favor compensatory heterochromatinization at susceptible loci such as Ighv.

### TADs and loops

At the finer level of 3D architecture – TADs and chromatin loops – we observed divergent dynamics between aging, S6-KO and CK-p25 neurons. In aged neurons, both TADs and loops became *more* pronounced, accompanied by elevated cohesin occupancy. These changes align with increased local compaction observed in aged neurons and with previous reports of enhanced local 3D contacts in senescent cells^74^. Thus, aging appears to preserve cohesin-mediated loops (even increasing average loop size), suggesting a compensatory tightening of local chromatin architecture. In contrast, S6-KO neurons displayed *weaker* TADs and reduced loop strength. CK-p25 neurons paralleled S6-KO, showing weakened TADs and loop loss, and the prominence of TAD and loop erosion again increased with the severity of the condition (S6-KO → Step 1 → Step 2).

Importantly, the loops lost (i.e., down-folded) in aging, S6-KO and CK-p25 neurons were predominantly those lacking gene regulatory elements at their anchors – i.e. structural or architectural loops thought to be formed by CTCF-cohesin binding between distant loci. Conversely, loops that contained promoter or enhancer elements at their anchors (putative regulatory loops) tended to be preserved (in S6-KO and CK-p25) or even strengthened (in aging). These observations suggest a model in which the 3D genome remodeling skews toward loss of structural scaffolds (likely due to weakening of CTCF/cohesin-mediated constraints^19,43,73,75^, while short-range gene regulatory interactions regulated by TFs persist, suggesting that chromatin stability is at least partially maintained by the activity of these genes. Our combined data hints that DNA damage and deficient chromatin repair may underlie the collapse of architectural loops in neurodegeneration.

### Transcriptomic changes

In parallel with the 3D genome changes, we compared transcriptomic alterations and found notable overlaps as well as condition-specific signatures. There were opposite gene expression changes between S6-KO and aging, but S6-KO and CK-p25 (Step1 and Step2) neurons showed a significant overlap of DEGs, with expression differences intensifying along the neurodegenerative trajectory (S6-KO → Step 1 → Step 2), mirroring the trends observed in chromatin architecture – erosion of TADs and loops, strengthening of B-B compartment interactions, etc As suggested before, it is possible that TF networks can change and maintain 3D structure, and our results showed that a shared set of such TFs in S6-KO and CK-p25. Among them, REST emerged as a central regulator associated with increased chromatin interactions and up-regulated DEGs in SIRT6-deficient neurons. Remarkably, this fits our previous results showing that SIRT6 interacts with REST and the Polycomb component EZH2 to modulate epigenetic silencing^61^.

Other TFs showing coordinated chromatin and transcriptional shifts included TBP, TP53, MYC, and E2F1, regulators of cell cycle re-entry, apoptosis, and stress response^65,66,67,68^. Their enrichment in both SIRT6-deficient and neurodegenerative contexts points to a convergent activation of stress-responsive transcriptional programs. Additionally, members of the Polycomb complex (SUZ12, EZH2, and BMI1) were consistently associated with regions of altered chromatin interactions and upregulated genes across SIRT6 deficiency, aging, and neurodegeneration. This pattern supports a broader role of Polycomb-mediated repression and chromatin compaction in shaping neuronal functions.

### Limitations of the study

The main limitation of this study lies in the potential divergence of chromatin architectural outcomes between humans and mouse models due to species-specific variations. This should be considered when extrapolating the findings to human neurodegenerative processes.

In this study, we obtained Hi-C data from neurons sorted by NeuN staining, complementing previously published datasets in which neurons were separated by both H2AX and NeuN markers. Although applying identical sorting protocols could further improve the results, we observed clear similarities and gradual changes across these neurodegenerative models.

We also used shRNA-mediated knockdown in primary cortical cultures (reducing SIRT6 levels by approximately 50%) and compared the resulting transcriptomes to RNA-seq data from cortical neurons in the CK-p25 system, characterized by an aggressive accumulation of DNA double-strand breaks and subsequent genomic instability. Despite the contrasting severity of these models, the mild shSIRT6 perturbation produced transcriptomic alterations that share similarities with those observed in the highly aggressive p25-induced phases.

## Conclusions

In summary, chromatin and transcriptional outcomes of our S6-KO model closely parallel those of the more severe CK-p25 system, implying shared pathogenic mechanisms and highlighting SIRT6 as a critical stabilizer of chromatin structure. SIRT6 deficiency disrupts neuronal 3D genome organization, weakens loop and TAD integrity, and activates stress-responsive transcriptional networks, together resembling features of neurodegeneration. Normal aging, in contrast, produces more contained and compensatory chromatin changes. These findings support a model in which the aging brain remains resilient, while loss of SIRT6-mediated chromatin maintenance precipitates architectural collapse and transcriptional instability – a hallmark of Alzheimer’s disease. Strategies to restore or mimic SIRT6 activity may therefore hold promise in preserving chromatin integrity and delaying neurodegenerative decline.

## Data availability

Sequencing data is under uploading to GEO (https://www.ncbi.nlm.nih.gov/geo/). Raw data of the previously published Hi-C and RNA-seq experiments for Step1, Step2 and Control neurons from CK-p25 mice is deposited under GSE227445 (Hi-C) and GSE174265 (bulk RNA-seq) GEO accessions. Public snRNA-seq data of Old and Juvenile mouse brains was downloaded from CZ CELLxGENE website: https://cellxgene.cziscience.com/collections/31937775-06024e52-a799-b6acdd2bac2e. The code used for the analyses in this study is available on GitHub: https://github.com/SIRT6/Sirt6_hic_paper.

## Acknowledgements

The RNA-seq data analysis was supported by the Russian Science Foundation (grant number 25-71-20017 to EK). The European Research Council (ERC) under the European Union’s Horizon 2020 research and innovation program (grant agreement No 849029 to D.T.), the David and Inez Myers foundation (D.T.), the Israeli Science Foundation 422/23 (D.T).

## Author contributions

E.Ka. and D.S. performed bioinformatic analysis and wrote the paper, E.E. performed Hi-C and microscopic analysis, A.T. performed microscopic analysis, A.G. conducted Hi-C experiments, D.K. performed bioinformatic analysis, S.K. performed animal experiments and RNA-seq, E.Kh. guided bioinformatic analysis, wrote the paper and planned the project, D.T. planned experiments, advised on experiments and results, and wrote the paper.

## Notes

### Competing Interest Statement

The authors have declared no competing interest.

