## Supplementary figures and images for "The Divergent 3D Genome Landscapes of Aging and Neurodegenerative mouse models"

### Supplementary fig 1

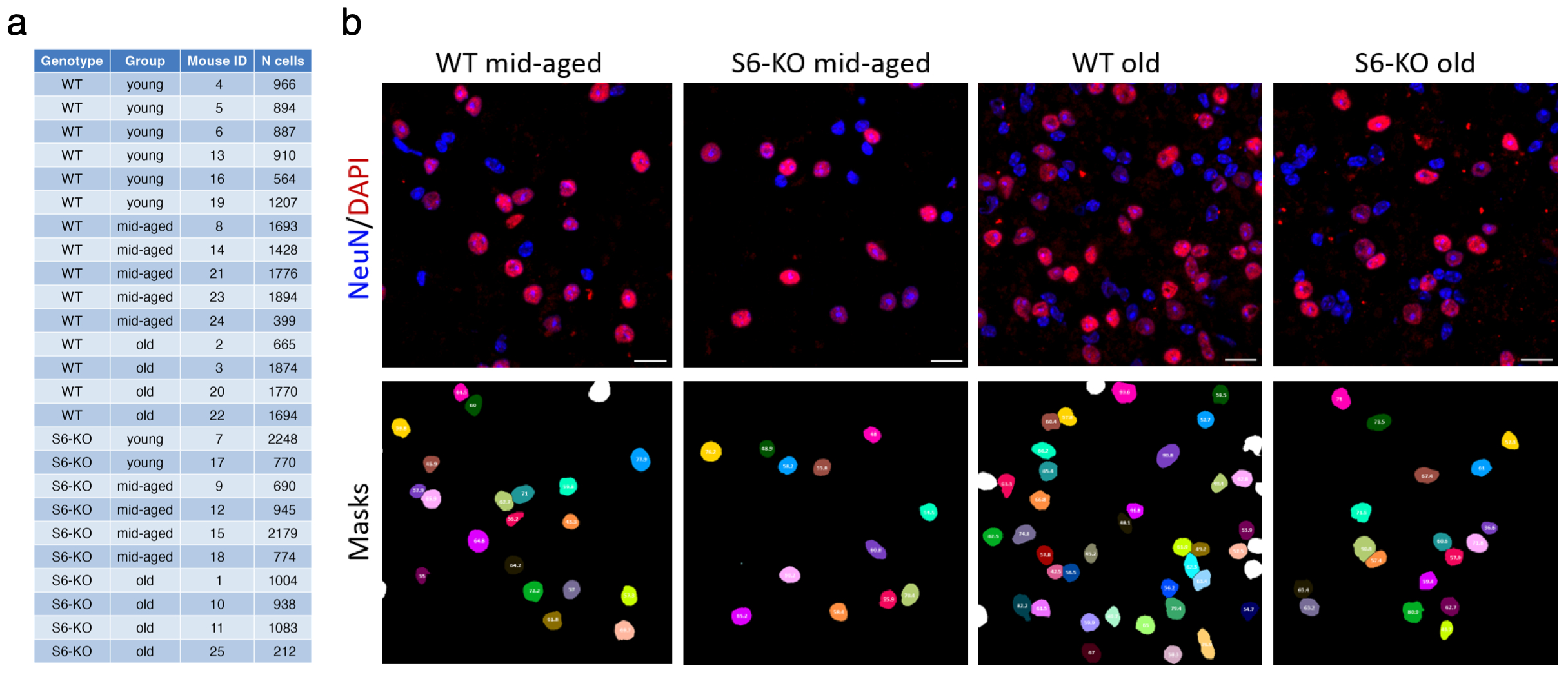

### Supplementary fig 2

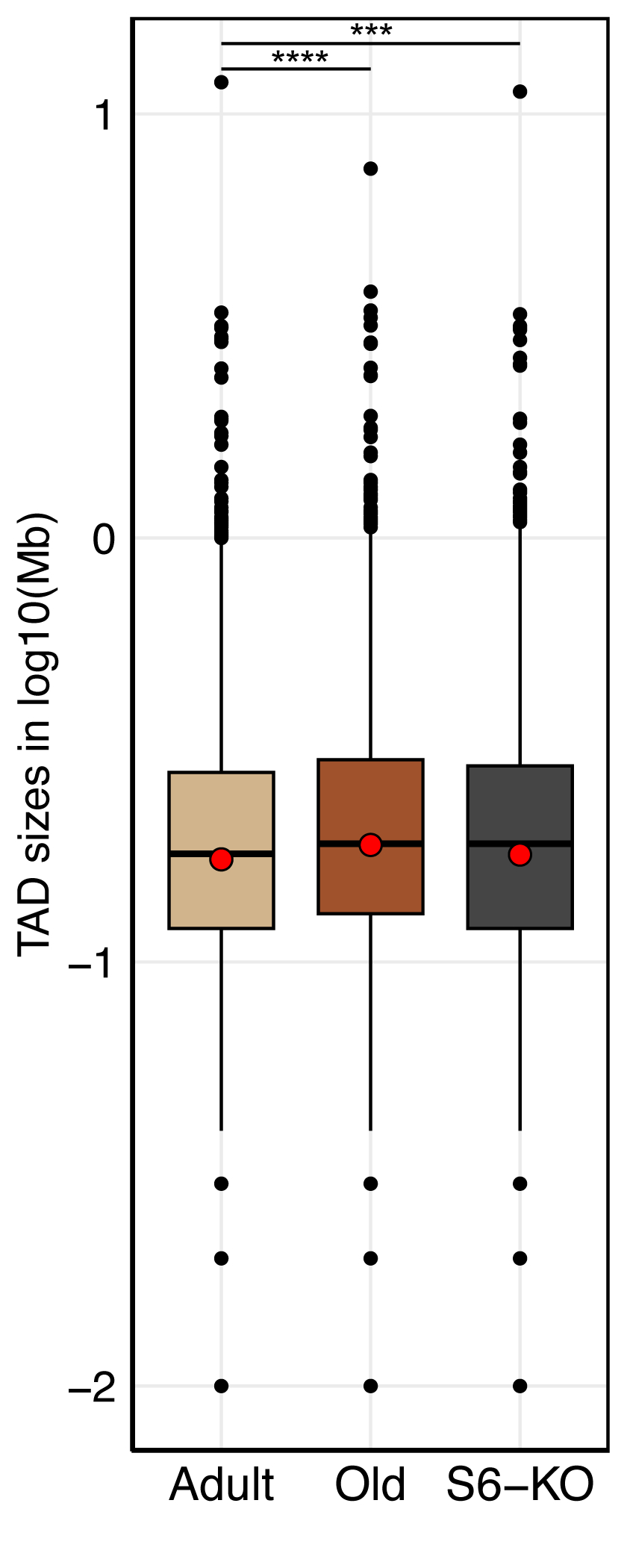

### Supplementary fig 3

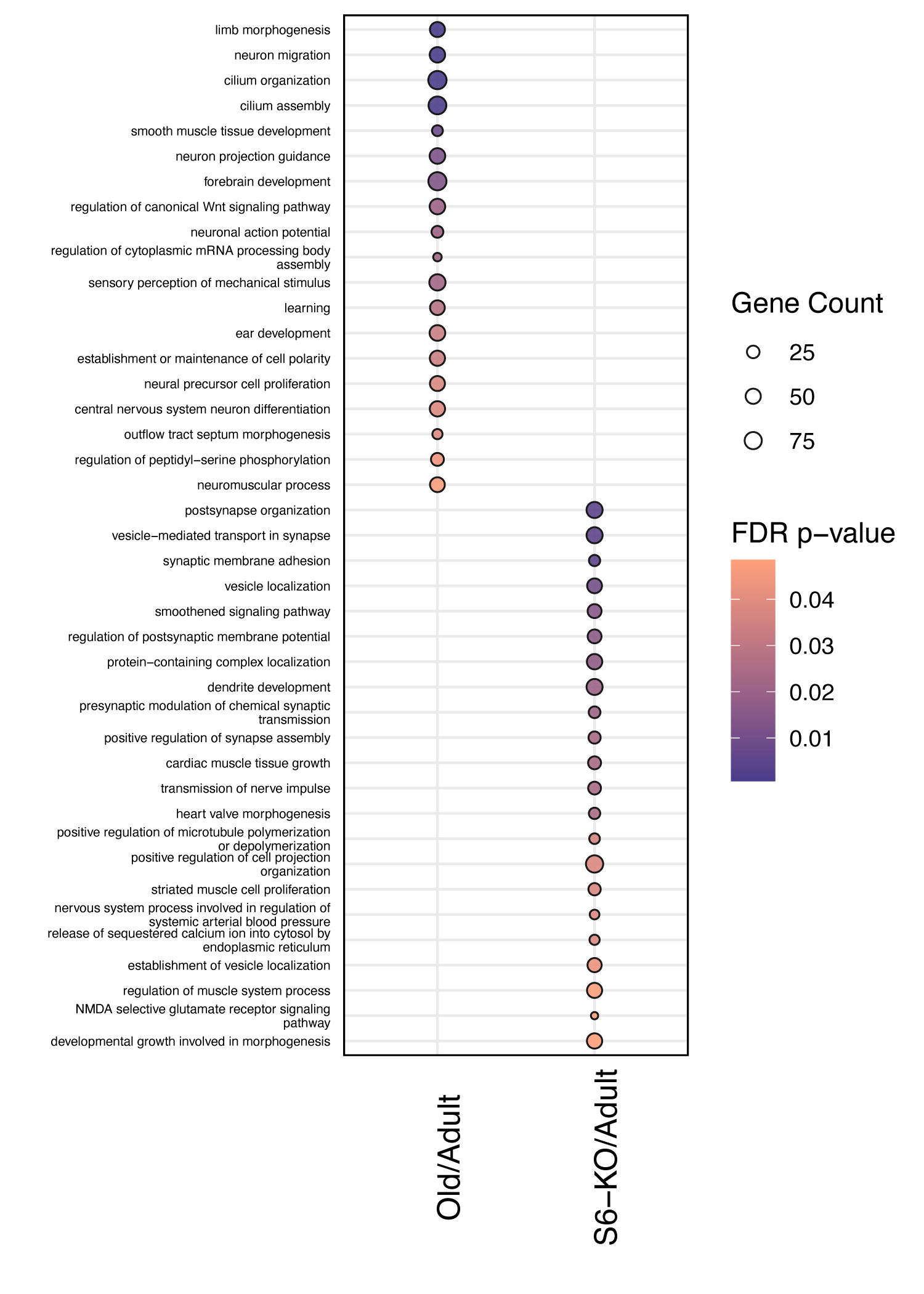

### Supplementary fig 4

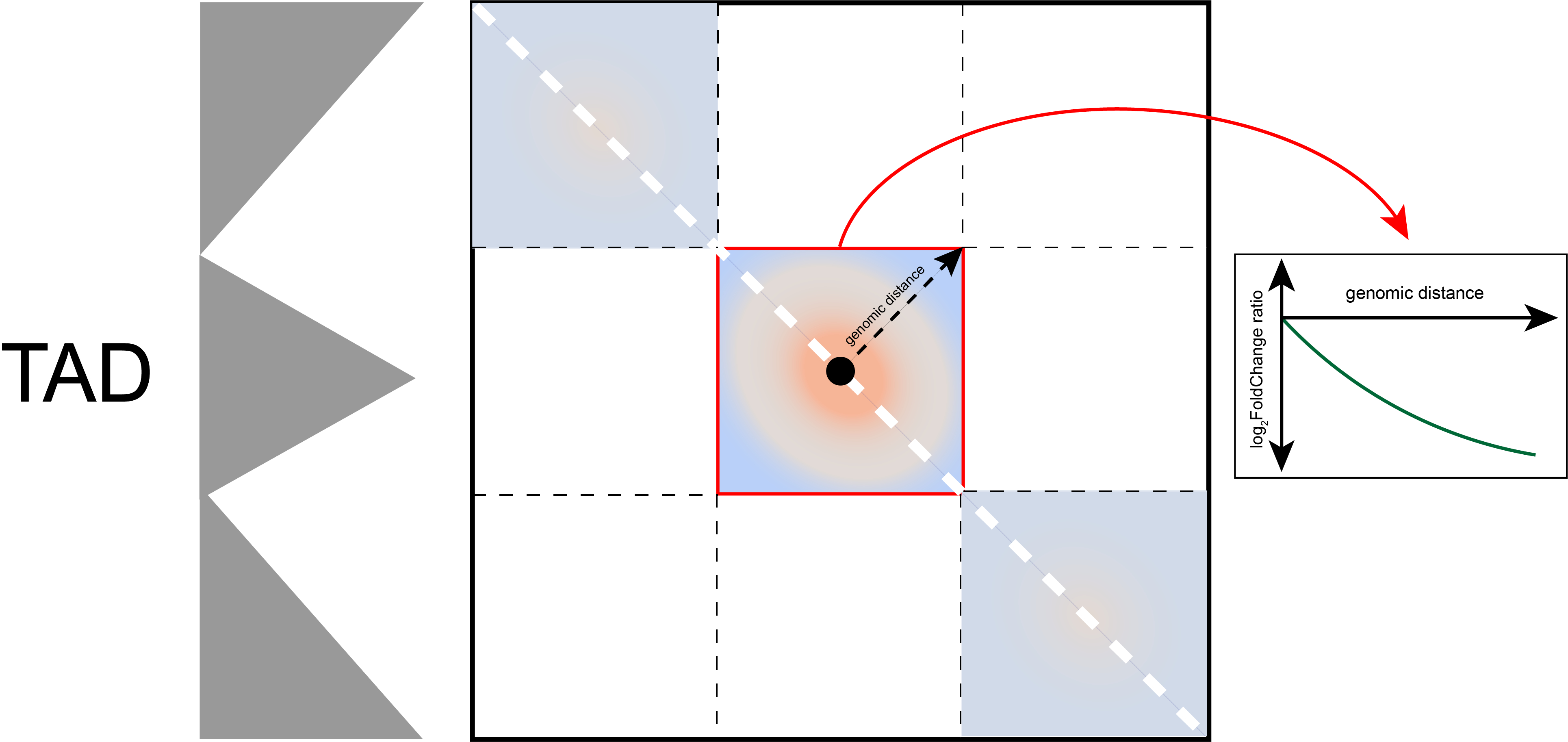

### Supplementary fig 5

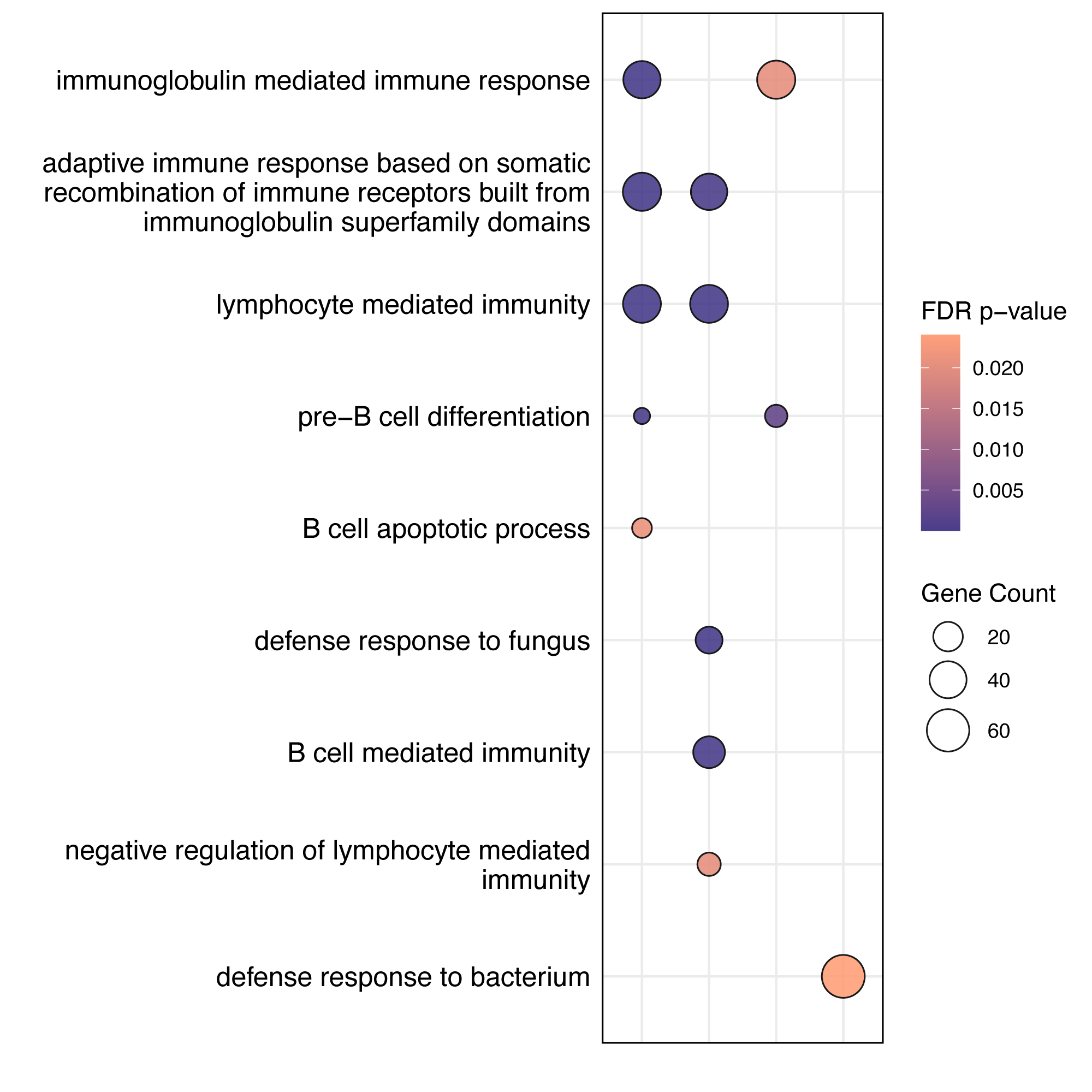

### Supplementary fig 6

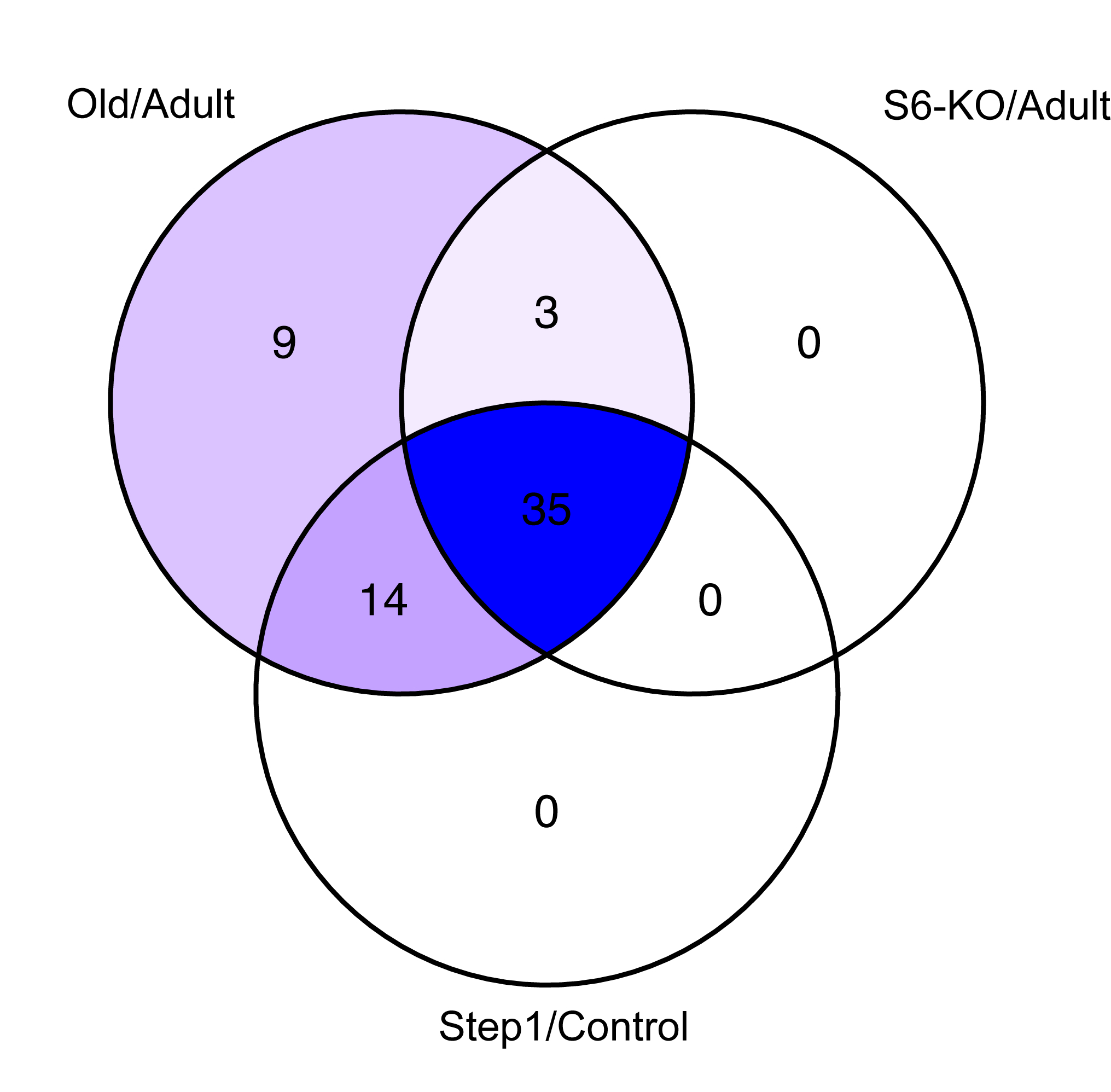

### Supplementary fig 7

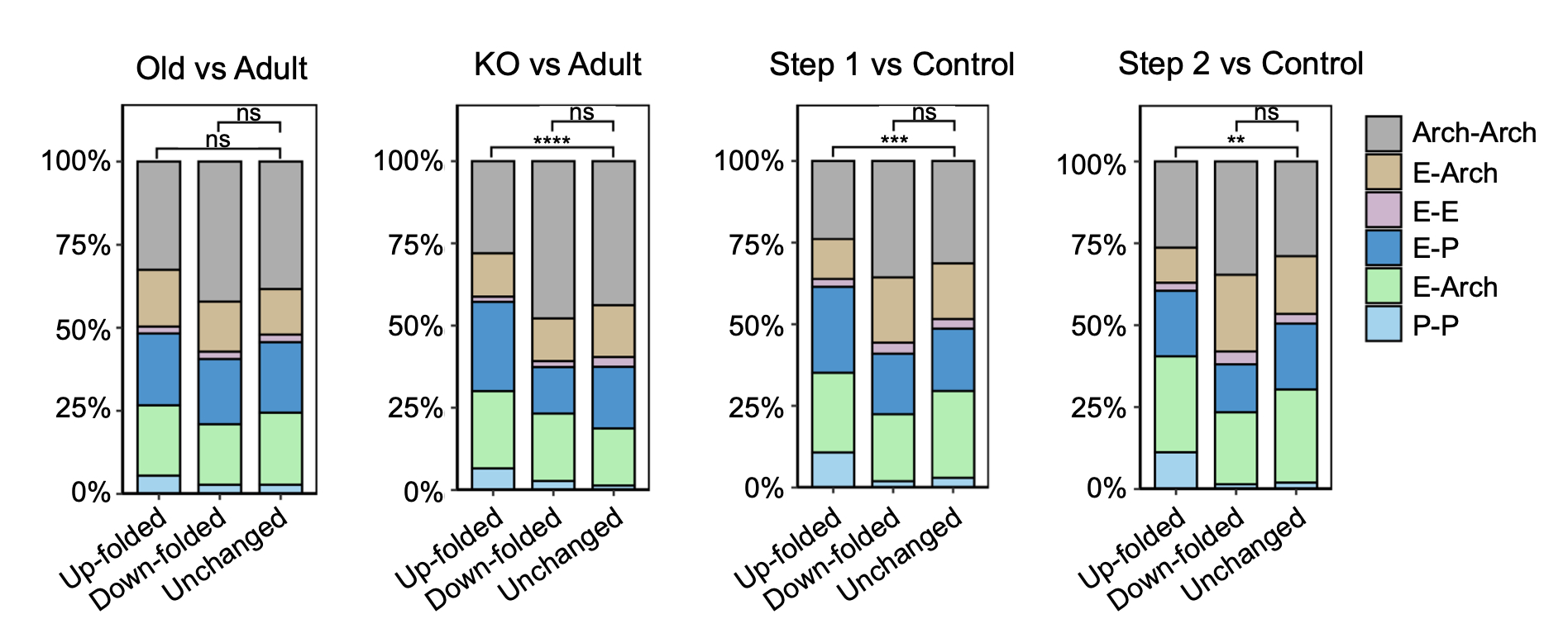

### Supplementary fig 8

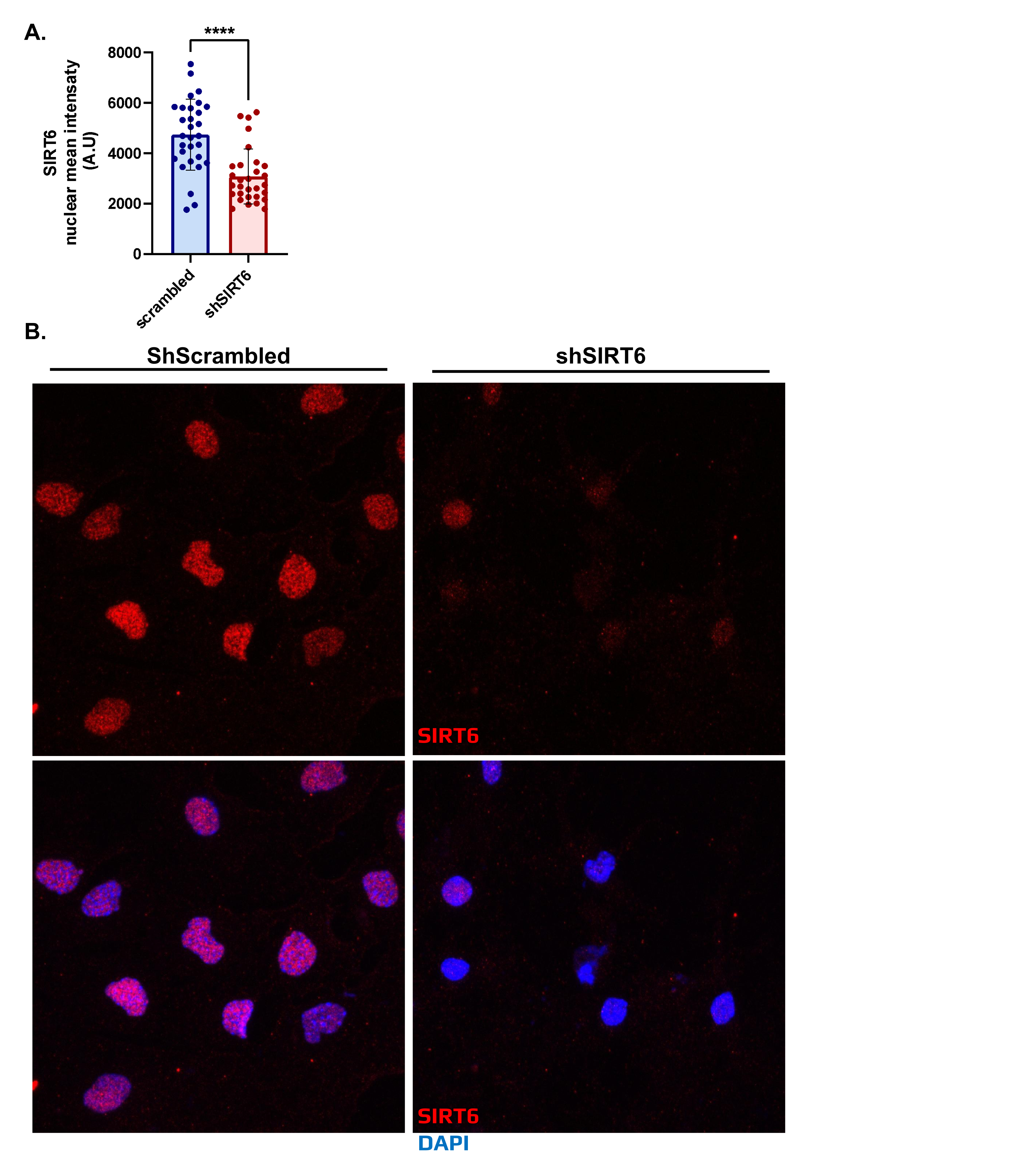

### Supplementary fig 9

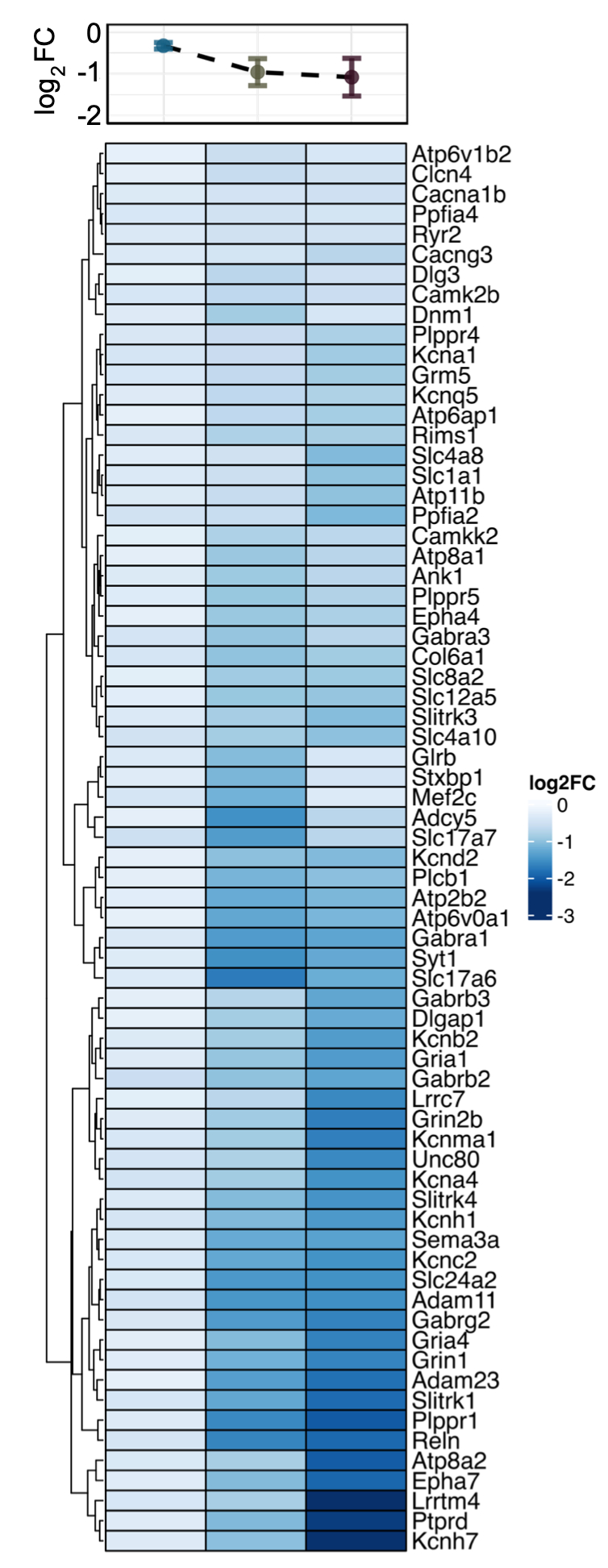
